# CDKN1A (p21^Cip/Waf1^) stabilizes Cyclin D3 by inhibiting its phosphorylation-dependent nuclear export following butyrate treatment

**DOI:** 10.1101/2025.06.19.660543

**Authors:** Mohammed Ala Ghrib, Guy W. Dayhoff, Cherif Chahtour, Carine Rodrigues-Machado, Abdelrahaman Youssef, Giorgio Schillaci, Guillaume Gautreau, Céline Henry, Vladimir N. Uversky, Pascal Rigolet, Hervé M. Blottière, Jean-Marc Lelièvre

## Abstract

Butyrate-mediated inhibition of cell proliferation is part of the preventive role of dietary fiber against colorectal cancer (CRC). This effect notably involves the cyclin-dependent kinase inhibitor CDKN1A (p21^Cip/Waf1^) in human intestinal cells, yet the underlying molecular mechanisms remain incompletely understood. Previously, we observed a paradoxical increase in cyclin D3 (CCND3)—but not cyclin D1—levels upon butyrate exposure. Here, we demonstrate that the butyrate-induced accumulation of CCND3 protein results both from mRNA increase and a CDKN1A-dependent protein stabilization, specifically extending its nuclear half-life. Proteomic analyses of CCND3 co-immunoprecipitates identified complexes involving CDKN1A, CDK4, CDK6, and the CRC-associated kinase CDK5, particularly enriched in butyrate-treated cells. Phosphorylation at a conserved Thr residue, crucial for CCND nuclear export and subsequent proteasomal degradation, was notably reduced following butyrate treatment and inversely correlated with CDKN1A expression levels. Structural modeling based on AlphaFold2, complemented by molecular dynamics simulations, revealed possible differential interactions between CDKN1A and cyclins D1 and D3, predicting that CCND3-Thr283 becomes structurally buried upon CDKN1A binding, limiting its phosphorylation. Our findings provide novel mechanistic insights into how CDKN1A might regulate CCND3 stability, highlighting previously unexplored roles of cyclin D3-containing complexes in cell cycle arrest induced by butyrate.

## 1. Introduction

Numerous studies have confirmed early findings that butyrate exposure in cell lines is toxic at high concentrations but inhibits cancer cell proliferation at lower doses [1]. This short-chain fatty acid (SCFA) is a major end-product of dietary fiber fermentation in the colon. Epidemiological and experimental data strongly suggest that dietary fibers prevent colorectal cancer (CRC) largely due to microbial production and action of butyrate in the colon [2–4].

Both butyrate and more specific inhibitors of histone/lysine deacetylases (H/KDACi) induce cell cycle arrest, particularly at the G1 phase, in a variety of cell lines and tumor models [5–8]. This arrest is frequently associated with cell differentiation or apoptosis [9]. Current evidence indicates that butyrate and H/KDACi treatments block the G1/S transition by decreasing CDK2 expression and activity while simultaneously enhancing the expression of CDK inhibitors CDKN1A, CDKN1B (p27), and CDKN1C (p57) [7, 8, 10–12]. Indeed, in HCT116 cells, this G1 arrest induced by butyrate and H/KDACi is CDKN1A-dependent yet independent of p53 [7, 13]. Consistently, elevated expression of CDKN1A alone can induce G1 or G2 arrest independent of p53 [14].

Previously, we demonstrated that treatment with butyrate or trichostatin A (TSA), a specific HDAC inhibitor, increases cyclin D3 (CCND3) protein levels in several intestinal cell types, including HCT116 cells [8]. Similar CCND3 induction was also observed in HCT116 cells exposed to the HDAC inhibitor apicidin [15], and in other contexts unrelated to butyrate. Interestingly, this increase appears paradoxical because cyclin D proteins (CCNDs) are well-established drivers of cellular proliferation, transformation, and tumor maintenance [16–19]. However, CCND functions extend beyond CDK activation alone [19–22] and their role is not always clear yet. Morevover, in some cases, the expression of the three type D cyclins may be differently regulated. suggesting that different signaling pathways and/or subtle structural differences may have evolved to respond to different challenges. For instance, CCND3 is abundant in quiescent cells in several human tissues. It is the case in the normal intestinal epithelium, while its expression is notably lost in colon adenomas [23]. CCND3 also plays crucial roles in skeletal muscle differentiation and gene expression [24, 25].

The expression and localization of type D cyclins are tightly regulated, as exemplified by CCND1 [19]. Newly synthesized CCND proteins must bind to CDKs to enter the nucleus [26, 27]. Conversely, nuclear export—necessary for cytoplasmic proteasomal degradation—is typically mediated by phosphorylation of conserved threonine residues within their C-terminal PEST domains [16, 28, 29]. Mutations preventing phosphorylation or loss of these residues often result in nuclear retention and are associated with certain cancers, as seen for CCND1 [28] and, more recently, CCND3 [30, 31].

Intriguingly, in two non-intestinal cell lines, CDKN1A was found to inhibit CCND1 nuclear export by binding the Thr286-phosphorylated form of CCND1 in dividing cells [32]. More broadly, CDKN1A and related proteins (CDKN1B, CDKN1C) exhibit dual regulatory roles on CCND-CDK complexes. Although predominantly known as tumor suppressors through their inhibition of CDK1/2 activity [33], they also promote cell cycle progression, particularly in cancer-like contexts, by modulating cyclin D levels and facilitating cyclin D-CDK4/6 nuclear entry [12, 34–37]. Inversely, cyclin D1 can inhibit CDKN1A protein degradation in certain cell contexts without altering cell division [38]. Structurally, CDKN1A is highly intrinsically disordered, possessing distinct regions dedicated to binding CCND-CDK complexes and proliferating cell nuclear antigen (PCNA) [39, 40].

Given these observations, we investigated whether the increase in CCND3 induced by butyrate or HDACi molecules arises primarily from enhanced mRNA expression or reduced protein degradation. Specifically, we evaluated CCND3 stability in the presence of inhibitors, compared expression levels in CDKN1A-deficient cells, assessed subcellular localization following butyrate treatment in HCT116 cells, and tested interactions between CDKN1A and native or recombinant CCND3. To further elucidate the molecular mechanisms involved, we performed mass spectrometry on CCND3-associated protein complexes and conducted structural analyses and molecular dynamics simulations of CDK4-CCND1/3-CDKN1A complexes. Our findings offer novel insights into butyrate’s cellular effects and provide a structural model suggesting a testable mechanism through which CDKN1A stabilizes CCND3.

## 2. Material and methods

### 2.1. Chemicals and antibodies

Most chemicals were purchased from Merck Sigma-Aldrich except those mentioned below. Tris-buffered saline (TBS) and anti-rabbit antibodies coupled to FluoProbes 488 were purchased from Interchim (France). Acrylamide gradient gels were purchased from Bio-Rad France and Thermo Fisher Scientific. Hoechst 33342 and reagents for reverse transcription and PCR were purchased from Thermo Fisher Scientific. Secondary anti-mouse and anti-rabbit antibodies conjugated to horseradish peroxidase (HRP) were purchased from Abliance (France). The anti-cyclin D3 rabbit polyclonal antibodies used in this study were purchased from Santa Cruz Biotechnology (sc-182) and Proteintech (26755-1-AP). Rabbit monoclonal antibodies against CDKNA1, Thr283-phosphorylated Cyclin D3, Thr286-phosphorylated cyclin D1, HA (influenza hemagglutinin) tag, and Mcl-1 were purchased from Cell Signaling Technology (USA). Mouse monoclonal antibodies directed against GAPDH (Santa Cruz Biotechnology, USA) and β-actin (Progen Biotechnik, Germany) were also used.

### 2.2. Cell culture and treatments

CDKN1A (p21)-/-cells and their parental HCT116 cell line were provided by Prof. Vogelstein’s laboratory (Johns Hopkins University School of Medicine, Baltimore, USA). The parental HCT116 cell line was used to generate the clones expressing CCND3-HA. The HCT116 clones expressing CCND3-HA protein (with the HA tag GYPYDVPDYA added at the C terminus) were obtained after transfection with the plasmid pcDNA3-CCND3 [41], kindly provided by Dr. Doris Germain (Peter MacCallum Cancer Institute, Victoria, Australia), and subsequent isolation of antibiotic-resistant colonies. These clones, the CDKN1A-/-cell population, and the associated parental cell line were maintained in RPMI Medium 1640 + GlutaMax supplemented with 10,000 U Penicillin, 10 mg/ml Streptomycin, and 10% fetal calf serum (FCS), and were incubated in 5% CO2 at 37°C. They were cultured in the presence of 350 µg/mL G418, except when used for experimental treatments. This was particularly important because the CDKN1A-/-cell line appeared to be contaminated by parental-like cells (i.e., cells expressing CDKN1A protein) in all lots obtained from different sources (which all originated from the same original culture). We also used a parental HCT116 cell line purchased from the American Type Culture Collection (ATCC) for this work. These latter cells were cultured in McCoy 5A medium (10,000 U Penicillin, 10 mg/ml Streptomycin, 10% FCS) in 5% CO2 at 37°C, as recommended by the ATCC. The Caco-2 cell line was purchased from ATCC and cultured in DMEM (10,000 U Penicillin, 10 mg/ml Streptomycin, 20% FCS).

### 2.3. CRISPR/Cas9-mediated CCND3 C-terminal deletion in HCT116 cells

We used the KN2.0 kit from Origene, which includes a donor cassette for a non-homology mediated CRISPR knockout, following the manufacturer’s instructions (https://www.origene.com/products/gene-expression/crispr-cas9/knockout-kits). Colonies resistant to puromycin (1.5 µg/ml) were screened for the absence of full-length cyclin D3 by Western blotting of total protein extracts. The mutant used in this work was generated using the RNA guide (5′-AGCATTGGGGCTGCAGTGCA-3′) in the cyclin D3 (CCND3) gene, which modified the sequence downstream of Ala211 (Fig. S1, Supplementary Data).

### 2.4. Measurement of cell proliferation

We followed the protocol described by Vichai *et al*. [42], which is based on the staining of fixed cells with Sulforhodamine B (SRB), a dye that binds to cellular proteins. Each well of a 96-well plate received 8,000–10,000 cells. After 24 hours of culture (Day 1), the culture medium was replaced with fresh medium with or without butyrate, and the cells were incubated further. The medium was changed every 2 days. After the incubation period, the amount of released SRB was measured using a TECAN plate reader.

### 2.5 Flow cytometry and FACS

Flow cytometry was used to determine the distribution of cells in various phases of the cell cycle. Briefly, cells treated or not treated with butyrate were detached in the presence of trypsin, washed twice with ice-cold PBS, and then fixed in 70% ice-cold ethanol for at least 1 hour at – 20°C before analysis. The fixed cells were washed twice with PBS containing 1% BSA and resuspended in this solution supplemented with 7-AAD (1.5 µg/ml), RNase (0.1 mg/ml), and 0.05% Triton. After 1 hour at 37°C, cells were resuspended in PBS/0.5 mM EDTA and immediately analyzed on a FACSCalibur or Fortessa x20 (Becton Dickinson).

The cytometer parameters were adjusted to separate single cells from aggregates by measuring fluorescence peak height (FL3-H), width (FL3-W), and area (FL3-A) of DNA-bound 7-AAD. The percentages of cells in G1, S, and G2/M phases (FL3-H histogram) were determined using Flowing Software 2.5.1 (https://bioscience.fi/services/cell-imaging/flowing-software/), developed by Perttu Terho (Turku Bioscience Centre, Turku, Finland).

### 2.6. Preparation of Cell Lysates for Western Analysis

Cell lysates and Western blot analyses focusing on the effect of butyrate were prepared as previously described [43] using a low-detergent buffer [44] to extract total “soluble” proteins. Approximately 2.3 million cells were seeded onto each 60 mm tissue-culture dish. Butyrate treatment began the following day (≥ 24 hours after plating) and lasted for at least 16 hours. The extraction buffer was supplemented with protease and phosphatase inhibitor cocktails (PhosphoSTOP, Roche; Complete Mini, Sigma 11836153001). Samples were run on either in-house prepared or commercial 8–15% gradient Tris-Glycine SDS-PAGE gels (Bio-Rad). To monitor equal protein loading, we performed immunodetection using GAPDH or β-actin antibodies. After the signal was developed for each primary antibody, HRP activity coupled to the secondary antibody was neutralized by concentrated hydrogen peroxide to enable reprobing of the same blot with an additional antibody.

### 2.7. Extraction and Western analysis to monitor the effect of inhibitors

Treatments with Cycloheximide (CHX) and Mithramycin A (MTM) were conducted in 24-well plates. Each well received approximately 200,000 cells, and treatments began the following day. For simultaneous co-treatments, the inhibitors were added to fresh medium supplemented with 1.5 mM butyrate. To assess the effects of inhibitors after butyrate pretreatment, 1 µL of a 1000× stock solution of CHX or MG132 was added directly to the culture medium 16 hours after the start of the 1.5 mM butyrate treatment ([Butyrate]_init). After incubation for the indicated times, total proteins were recovered in either 1.5× Laemmli-based (1.5% LDS, 49.35 mM Tris-HCl pH 6.8, 30% glycerol, 50 mM TCEP; Bio-Rad) or NuPage LDS (Thermo Fisher Scientific) sample buffer. The extracts were then heated, centrifuged (as described previously [45]), and analyzed by SDS-PAGE or NuPage Bis-Tris gradient gels, respectively.

### 2.8. Quantification of Western signals

At least three independent Western blot analyses were quantified to generate graphs and perform statistical analyses. Band intensities were measured using ImageJ (Fiji, https://imagej.nih.gov/ij) with the Measure plugin, as well as using the iBright program (Thermo Fisher) with local background correction. The luminescence signals were normalized by dividing the intensity of the protein of interest by that of actin or GAPDH. To compare data from different Western analyses, we either normalized the values to 1 (where the highest signal on each gel was set to 1) or used the T0 signal (as shown in Fig. 2(d)) as the reference.

**Fig. 1:**
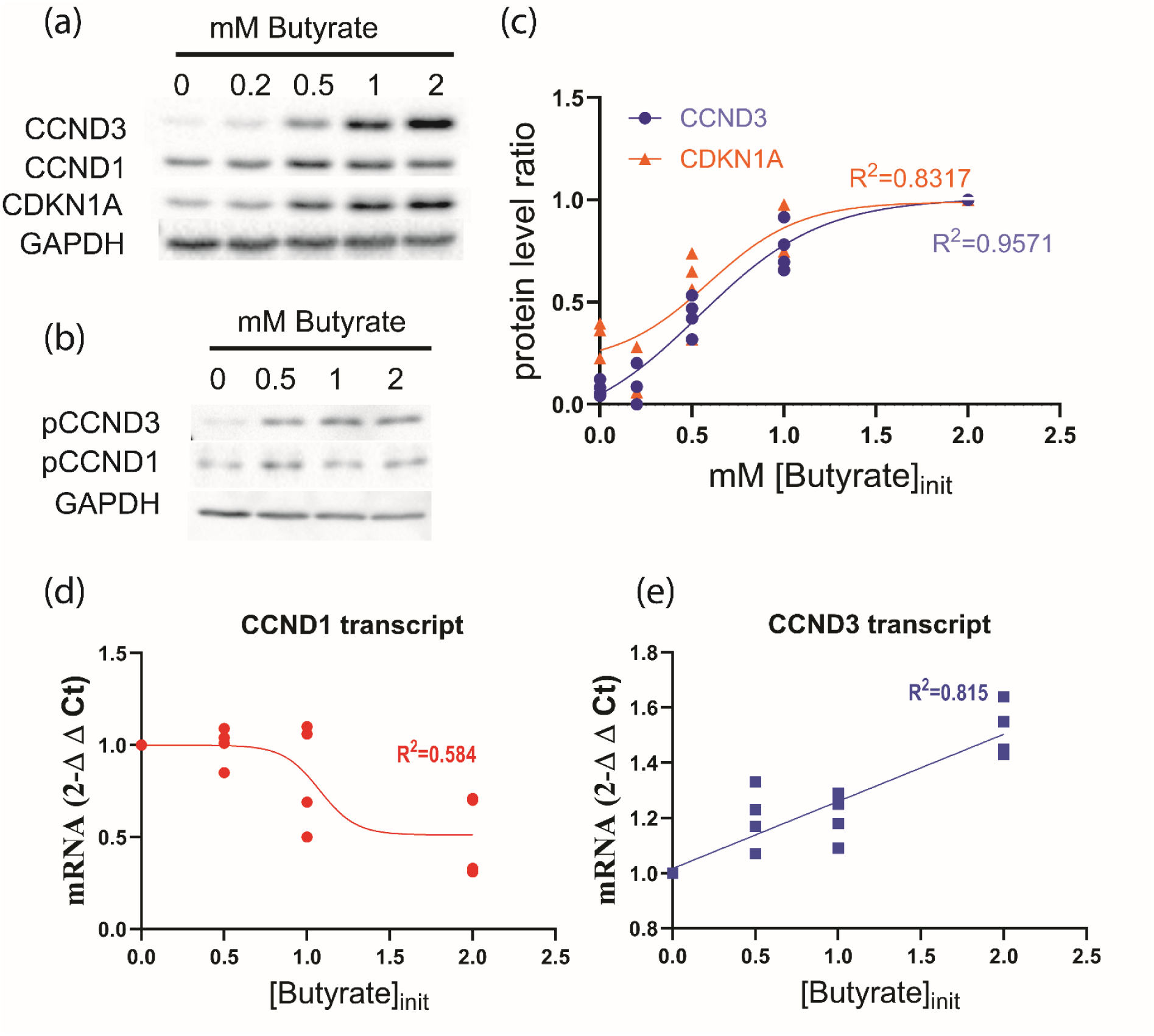
Dose response to butyrate for cyclins D1 (CCND1), D3 (CCND3) and the cyclin-dependent-kinase (CDK) inhibitor CDKN1A protein, and CCNDs mRNA levels in HCT116 cells. Total “soluble” proteins were extracted from cells cultured during 16 hours in a medium containing initially different butyrate concentrations. (a) Representative western analysis; (b) quantitative analysis of the normalized western signal (level) ratios derived from 3 independent experiments; (c) Western analysis of phosphorylated cyclins D (same extracts as used in (a)). In (c), the sigmoidal fit for both CCND3 (R^2^= 0.9571) and CDKN1A (R^2^=0.8317) protein level ratios are shown. (d)(e) Dose response to butyrate treatment for transcript levels of CCND1 and CCND3 genes measured in HCT116 by RT-qPCR. Measurements issued from three different experiments were plotted. The graphical representation indicated a linear increase in CCND3 mRNA (P< 0.0001) while CCND1 mRNA level decreased (P=0.001).

**Fig. 2:**
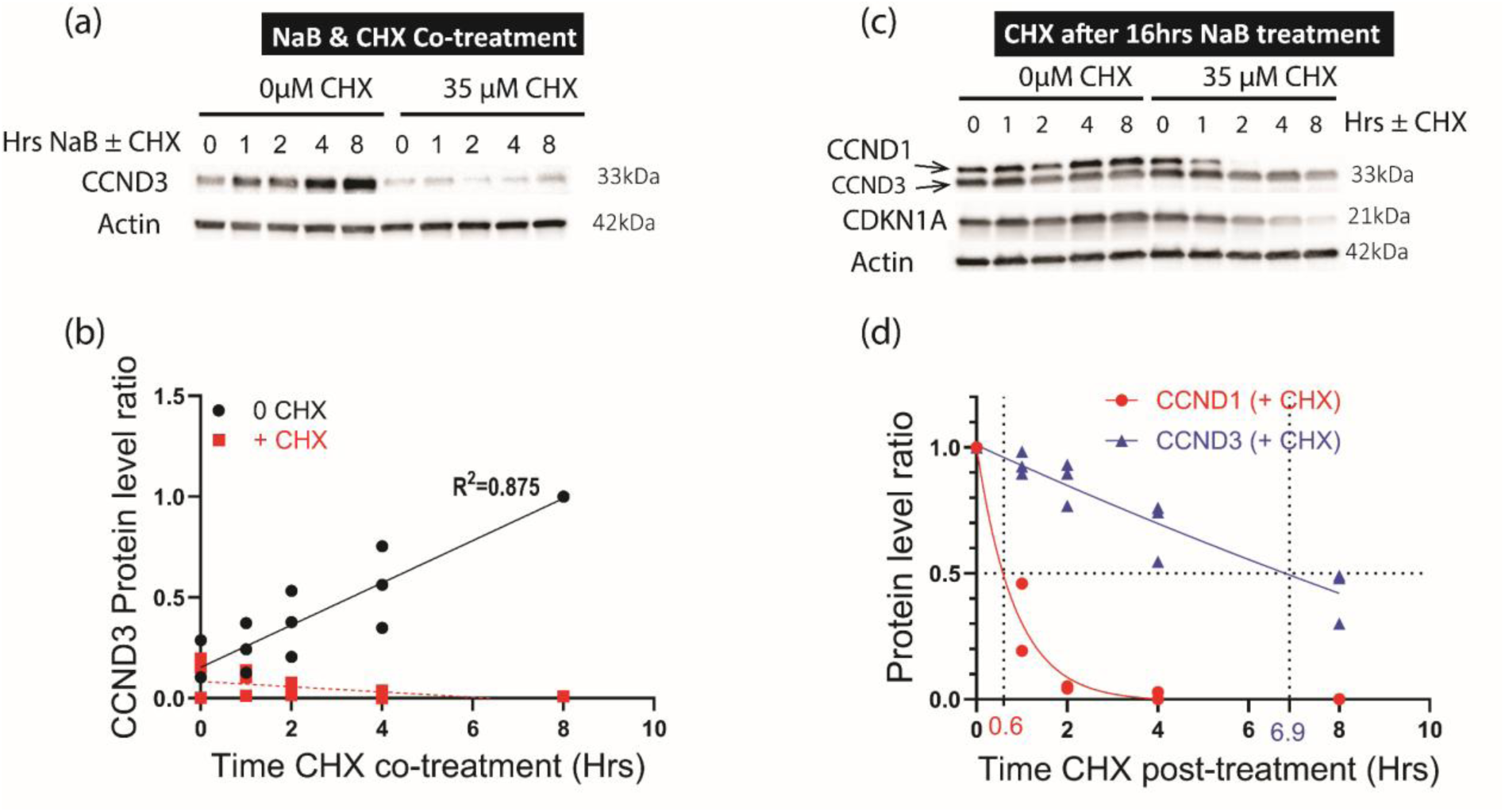
The butyrate-dependent increase in CCND3 protein level in HCT116 cells required both protein neosynthesis and stabilization. (a)(b) Total proteins were recovered from cells that were cultured into a medium containing 1.5 mM butyrate with 36 µM CHX or DMSO during 0 to 8 hours. (a) Images of a representative western analysis. (b) Graphical representation aggregating 3 independant analyses. The best fit for the protein level ratio was obtained with a linear regression both in the absence of inhibitor (R^2^=0.875; P<0.0001) and in its presence (R^2^=0.275; P=0.0446). <u>(c)(d)</u> CCND3 protein displayed a longer half-life than CCND1 in butyrate-treated cells. Cells were first cultured in a medium containing initially 1.5 mM butyrate during 16 hours, and then exposed to 36µM CHX or DMSO only directly added to the medium for 0 to 8 hours. Proteins were extracted as in Fig. 2. (a) Representative western analysis for CCND3, CCND1 and CDKN1A. (b) Graphical representation of the function displaying the decreased protein level ratio for CCND1 (R^2^=0.972) and CCND3 (R^2^=0.900) in cells exposed to CHX.

### 2.9. Reverse transcription and quantitative PCR (RT-qPCR)

Cells were harvested after 16–18 hours of butyrate treatment as described above. Total RNA was isolated from 1–3 million HCT116 cells using RNeasy Mini spin columns (Qiagen, Germany), and the RNA concentration was determined on a NanoDrop spectrophotometer (Fisher Scientific, France). Two micrograms of RNA from each sample were mixed with the reverse transcriptase mix (High-Capacity cDNA Reverse Transcription Kit, Thermo Fisher Scientific, USA) supplemented with oligo-dT primers. The reactions were performed at 25°C for 10 min, 37°C for 120 min, and 85°C for 5 min. For quantitative PCR, the optimal cDNA dilution was first determined to ensure linear amplification. qPCRs were run in triplicate with TaqMan Master Mix and TaqMan primers (Thermo Fisher Scientific, USA) in a 20 µL final volume using a StepOne Plus system (Applied Biosystems, USA). The results were analyzed with QuantStudio software (Qiagen, Germany). Values were calculated as relative-fold differences using β-actin as the reference gene, and the 2^−ΔΔCT method was used for expression. The analysis was performed three times from three independent cell treatments. All experiments were performed according to the manufacturers’ recommendations.

### 2.10. Immunofluorescence

Cells were fixed for 20 minutes at room temperature in 4% formaldehyde, rinsed three times with PBS, and permeabilized in 0.1% Triton X-100 for 10 minutes. Nonspecific binding sites were blocked in TBS containing 0.05% Tween 20 (TBST) and 2% bovine serum albumin (BSA). Primary antibodies were applied overnight at 4°C. After several washes with TBST, cells were incubated for 1 hour at room temperature with anti-rabbit IgG conjugated to a fluorochrome. Following three additional washes in TBST, the DNA was stained for 10 minutes in TBS containing 1.5 µg/ml Hoechst 33342. Samples were then rinsed, mounted with Vectashield, and observed with a MetaXpress HCS microscope (Molecular Devices Inc.) under confocal mode (pinholes/slits) using a 20× objective.

### 2.11. Co-immunoprecipitations (Co-IP)

Cell proteins were extracted and quantified in Lysis Buffer (#9803; Cell Signaling Technology) supplemented with protease (cOmplete Mini, Sigma 11836153001) and phosphatase (PhosphoSTOP, Roche 04906837001) inhibitor cocktails, according to the manufacturer’s instructions. Primary antibodies (anti-HA, 26181-2, Pierce Thermo Scientific) were directly immobilized on agarose beads, while Protein A–conjugated magnetic beads (Cell Signaling Technology, USA) were used to pull down complexes with non-conjugated antibodies (anti-CCND3 and anti-CDKN1A).

We largely followed the Pierce HA tag IP/Co-IP agarose kit (Thermo Scientific) protocol. Briefly, 800 µg of extracted protein was mixed with primary antibodies or agarose beads and incubated overnight at 4°C with rotation. Immune complexes bound to HA-agarose beads were washed and purified using the provided columns or a magnetic rack. For samples with anti-CCND3 or anti-CDKN1A, Protein A–conjugated magnetic beads were added and incubated at room temperature for an additional hour prior to washing. After three washes in lysis buffer, beads were resuspended in 20–40 µL of 3× SDS or LDS loading buffer, supplemented with 50 mM TCEP, and heated for 10 minutes at 70°C before SDS-PAGE.

### 2.12. LC-MS analysis

Protein extracts were briefly separated (∼3 min) by 1D gel electrophoresis (NuPAGE 4–12% Bis–Tris Gel). After excising the gel bands, proteins were reduced with DTT, alkylated with iodoacetamide, and digested with 50 ng of trypsin. The resulting peptides were recovered, dried under vacuum, and resuspended in 40 µL LC-MS loading buffer (0.1% trifluoroacetic acid, 2% acetonitrile).

Peptide samples were analyzed using a timsTOF Pro mass spectrometer coupled to a nanoElute liquid chromatography system (Bruker Daltonik GmbH, Germany). Peptides were injected onto an Aurora C18 column (75 µm i.d. × 250 mm, 1.6 µm, 120 Å pore size; Ion Opticks, Australia) using a one-column separation method. Buffer A was 0.1% formic acid in 2% acetonitrile, and Buffer B was 0.1% formic acid in 100% acetonitrile. A 40-minute gradient was applied at 200 nl/min as follows: 2% to 5% B over 1 min, 5% to 13% B over 18 min, 13% to 19% B over 7 min, and 19% to 22% B over 4 min, followed by a high-organic wash at 95% B for 7 min. The column temperature was 50°C.

Peptides were introduced via a CaptiveSpray nanoelectrospray ion source (Bruker Daltonik GmbH) at 1.6 kV, an ion source temperature of 180°C, with a dry gas flow of 3 L/min. Data-dependent PASEF (dda-PASEF) was used, with 6 PASEF ramps targeting precursors of charge 0–5 over a 1.30-s cycle. Each ramp was 180 ms, and ion mobility was calibrated in the 0.6–1.3 V·s cm^⁻²^ range using Agilent Tune mix. The mobility scan range was 0.7–1.10 V·s cm⁻², with collision energy increasing linearly from 20 eV at 0.60 V·s cm^⁻²^ to 59 eV at 1.60 V·s cm⁻². MS and MS/MS spectra were acquired from 100–1700 m/z. The TIMS accumulation time was 180 ms, with a target precursor intensity of 14,000 arbitrary units (au) and a threshold of 1000 au. Absolute intensity thresholds were set to 10 for mass spectra peak detection and 5000 for mobilogram peak detection.

### 2.13. Proteins identification

Database searches were conducted against a FASTA database derived from the Homo sapiens genome (48,434 entries) plus anti-HA sequence proteins, along with a list of common contaminants. We used i2MassChroQ (version 0.6.3, http://pappso.inrae.fr/) allowing for one missed cleavage. Cysteine residues were set as carboxyamidomethylated (fixed), and methionine oxidation was set as a possible modification. A precursor and fragment mass tolerance of 5 ppm was applied. Identified proteins were filtered and grouped using i2MassChroQ with the following criteria: peptide E-value < 0.01, protein log(E-value) < –4, and a minimum of two identified peptides per protein.

### 2.14. Analysis of the proteomic data

Three independent replicates of each CCND3-HA co-immunoprecipitation sample (from L3 and P3 extracts with or without butyrate treatment) were analyzed by LC-MS, yielding several hundred proteins in each run. To identify the proteins most likely to interact with CCND3-HA and eliminate false positives, a rigorous screening procedure was applied (The corresponding R script and data are available at the following link: “ https://forgemia.inra.fr/metagenopolis/cyclin_d3/<u>”</u>). Briefly, after removing all ribosomal and histone proteins, the polypeptides detected in the CCND3-HA IP fraction from two parental butyrate-treated HCT116 cell samples (negative controls “CTRL1” and “CTRL3”) were also retrieved from the L3 and P3 samples. We then selected proteins found exclusively in the six L3/P3 replicates without butyrate (L3/P3_noButyrate) or exclusively in the six replicates with butyrate (L3/P3_PlusButyrate). Finally, to compare the abundance of proteins identified in both no-butyrate and butyrate conditions, each protein’s PAI value was normalized by dividing it by the PAI of CCND3-HA in the same sample.

### 2.15. Disorder analysis

Per-residue disorder propensities and binding-induced folding propensities were evaluated using RIDAO (https://ridao.app) [46]. RIDAO yields predicted disorder propensities from PONDR VL-XT [47], PONDR VL3 [48], PONDR VSL2B [49], PONDR-FIT [50], IUPred Long, and IUPred Short [51, 52]. RIDAO also computes the mean disorder propensity, over all constituent predictors, for each residue. Furthermore, RIDAO yields binding-induced folding propensities from ANCHOR2 [53]. In all cases, propensities span the range 0-1 with values greater than or equal to 0.5 indicating positive propensity for disorder or binding-induced folding.

### 2.16. Atomistic model generation

Complete atomistic models of CDKN1A-CCND1/3-CDK4 were constructed using the partially resolved structure of CDKN1A-CCND1-CDK4 (PDBID: 6P8H) [36] along with AlphaFold2 (AF2) [54] predicted structures for CDK4 (UniProtID: P11802), CDKN1A (UniProtID: P38936), CCND1 (UniProtID: P24385), and CCND3 (UniProtID: P30281). AF2 structures for each chain were taken directly from the AF2 Database (https://alphafold.ebi.ac.uk/). The complex was created by superimposing backbone atoms of individual AF2 structures onto the respective chains in the crystallographic complex using PyMol [55]. The CDKN1A-CCND3-CDK4 complex was generated by superimposing the backbone atoms of CCND3 onto CCND1.

### 2.17. Molecular dynamics (MD)

MD simulations were performed using GROMACS 2020.4 [56] with the Amber99SB-ILDN [57] force field. Two systems were simulated: CDKN1A-CCND1-CDK4 and CDKN1A-CCND3-CDK4, with initial structures obtained as previously described (see: Atomistic model generation). Prior to minimization, steric clashes between chains were resolved by selectively rotating peptide bonds in intrinsically disordered regions (IDRs) using Rodrigues’ rotation formula. Three rotations were applied per complex: 180° at CDK4 residues 24–25 (chain B, C-terminal side), 120° at CCND1/3 residues 6–7 (chain A, C-terminal side), and 120° at CDK4 residues 61–62 (chain B, N-terminal side).

Each system was solvated in a cubic box with 1 nm padding using the SPC/E water model. Na⁺ and Cl⁻ ions were added to neutralize the system and achieve a 100 mM ionic concentration. Periodic boundary conditions were applied in all directions. Energy minimization was performed before and after solvation with an emtol setting of 500 kJ/mol/nm. Equilibration was carried out under NVT conditions for 100 ps using the Berendsen thermostat [58] (τ = 0.1 ps, T = 298 K), followed by 1 ns of NPT equilibration with the Berendsen barostat (τ = 1.0 ps, P = 1.01325 bar).

Production simulations were run under NPT conditions with the Parrinello-Rahman barostat [59](τ = 1.0 ps, P = 1.01325 bar) and the V-rescale [60 thermostat (τ = 0.1 ps, T = 298 K). All bond lengths were constrained using the LINCS algorithm, and electrostatics were handled with Particle Mesh Ewald {Darden, 1993 #16755], (PME) using a real-space cutoff 1.0 nm. Van der Waals interactions were also treated with a 1.0 nm cutoff. Simulations were conducted for 300 ns with a 2 fs timestep, and snapshots were saved every 25 ps for trajectory analysis.

### 2.18. Statistical analysis

Statistical analyses were performed using GraphPad Prism 9 (GraphPad Software, San Diego, USA) and R. We employed Mann-Whitney and Kruskal-Wallis non-parametric tests, followed by all-pairwise multiple comparisons with the Holm correction. Differences were considered significant at p < 0.05. Graphs were generated with Prism 9 or SigmaPlot v11 (Systat Software, Inc., USA).

## 3. Results and Discussion

### 3.1. Dose-dependent regulation of Cyclin D proteins by butyrate

First, we evaluated the impact of a 16-hour treatment with increasing concentrations of butyrate on cyclin D protein levels in HCT116 cells. Consistent with previous studies [8], we observed a steady increase in cyclin D3 (CCND3) levels following butyrate exposure. In contrast, cyclin D1 (CCND1) levels remained unchanged or showed only a modest increase at 0.5 mM butyrate (Fig. 1(a)), similar to earlier observations in HT29 cells [8]. Cyclin D2 (CCND2) was undetectable by western blot analysis in HCT116 cells, irrespective of butyrate treatment, confirming previous reports [61].

Interestingly, the dose-dependent responses of CCND3 and CDKN1A proteins to butyrate exhibited similar patterns. Their EC50 values (0.54 mM for CCND3 and 0.59 mM for CDKN1A) closely matched the IC50 value previously reported for butyrate-mediated inhibition of proliferation [62]. Levels of both proteins peaked between 1.5 and 2 mM butyrate (Fig. 1(c)), beyond which cell death increased sharply (data not shown). Additionally, both CCND3 (3.2.section) and CDKN1A (data not shown) levels increased within 4 hours of treatment.

Next, we investigated whether phosphorylation of cyclins D1 and D3 was altered by butyrate treatment, as phosphorylation typically precedes degradation for type D cyclins [16, 63, 64]. Following butyrate treatment, phosphorylation at Thr283 of CCND3 and Thr286 of CCND1 was detectable, albeit at low levels (Fig. 1(b)). In untreated cells, phosphorylation was barely detectable. Notably, the phosphorylation of CCND3 increased steadily alongside total protein levels in response to butyrate concentration. For CCND1, phosphorylation peaked at 0.5 mM butyrate, mirroring the pattern seen for total CCND1 levels (Fig. 1(a), (b)). These findings suggest either enhanced kinase activity or simply greater detection sensitivity due to increased cyclin levels in butyrate-treated cells.

Unlike HCT116 cells, CCND2 was detectable in Caco-2 cells and, along with CCND3, increased upon butyrate treatment up to 5 mM (Fig. S2, Supplementary Data). A similar butyrate-dependent induction of CCND2 has recently been reported in breast cancer cells [65]. Notably, in contrast to HCT116 cells [66], Caco-2 cells display higher differentiation levels [67, 68], indicating that butyrate-induced accumulation of cyclin D proteins does not depend strictly on cellular differentiation status.

Finally, we evaluated CCND3 and CCND1 mRNA levels following a 16-hour butyrate treatment. CCND3 transcript levels increased significantly, though moderately (P<0.0001), consistent with earlier reports suggesting transcriptional regulation by butyrate [15, 69]. In contrast, CCND1 mRNA levels significantly declined (P=0.001), reaching approximately 50% of control at 2 mM butyrate (Fig. 1(d)), as previously documented [70].

### 3.2. Dual regulation of Cyclin D3 (CCND3) stability: synthesis and degradation

Next, we re-examined the posttranslational regulation of CCND3 protein levels in butyrate-treated cells. During the first 8 hours of butyrate treatment, the CCND3 protein level increased significantly (P<0.0001) in the absence of the translation inhibitor cycloheximide (CHX). In contrast, simultaneous treatment with butyrate and CHX resulted in a moderate but significant (P=0.044) decrease in CCND3 protein levels (Fig. 2(a), (b)). These observations indicated that new protein synthesis was a critical factor during the initial phase of CCND3 accumulation induced by butyrate. However, we could not confidently determine the half-life of CCND3 in these initial conditions, as the semi-logarithmic plot of CCND3 protein levels over time showed a poor linear fit (data not shown). This likely resulted from the low initial CCND3 protein levels and asynchronous cell populations.

We then assessed the stability of CCND3 protein after 16 hours of pre-treatment with butyrate (Fig. 2(b), (d)), a condition in which most cells are blocked in the G1 phase of the cell cycle (see below) and butyrate concentrations in the medium have decreased substantially [71]. In control cells (without CHX), levels of both CCND3 and CCND1 remained relatively stable over the subsequent 8 hours. Remarkably, when CHX was added after the 16-hour pre-treatment, CCND3 protein levels declined more slowly compared to CCND1. The decrease in CCND3 protein level followed a linear trend when plotted on a semi-logarithmic scale (Fig. S3, Supplementary Data), allowing us to estimate a half-life of approximately 6.1 hours (range: 4.9–8.1 hours) for CCND3 (Fig. 2(d)). In contrast, CCND1 exhibited a considerably shorter half-life, estimated at 0.59 hours (range: 0.43–0.91 hours). This value for CCND1 is slightly longer than previously reported in synchronized cycling cells [61], where CCND1 was preferentially degraded during the S phase (half-life: 11 min) compared to the G1 phase (half-life: 27 min) in HCT116 cells.

These results demonstrate that posttranslational stabilization of CCND3 specifically occurs in butyrate-treated cells, correlating with their synchronized arrest in the G1 phase of the cell cycle (further detailed below and previously shown in [10, 72]. Thus, the butyrate-induced elevation in CCND3 levels depends both on de novo protein synthesis and on a marked reduction in its degradation rate.

### 3.3. CDKN1A stabilizes CCND3 protein levels following butyrate treatment

Given the strong correlation between butyrate-induced CCND3 stabilization and increased CDKN1A expression, we directly investigated whether CDKN1A was required for CCND3 accumulation. Previous studies showed that butyrate treatment fails to induce G1 arrest in the absence of CDKN1A ([7] and unpublished data), and CDKN1A has been implicated in the stabilization and nuclear localization of cyclin D complexes.

We observed significantly reduced CCND3 protein levels in CDKN1A^-/-^ HCT116 cells following butyrate treatment, although a modest residual increase remained detectable (Fig. 3(a),(b)). This small increase likely resulted from contamination with parental-like cells expressing residual CDKN1A protein, as indicated by a concurrent minor rise in CDKN1A expression (Fig. 3(c); see Materials and Methods). CCND3 and CDKN1A protein levels in the butyrate-treated CDKN1A^-/-^ cell population were significantly lower (**P<0.001**) than in parental cells, strongly suggesting a stabilizing role for CDKN1A.

**Fig. 3:**
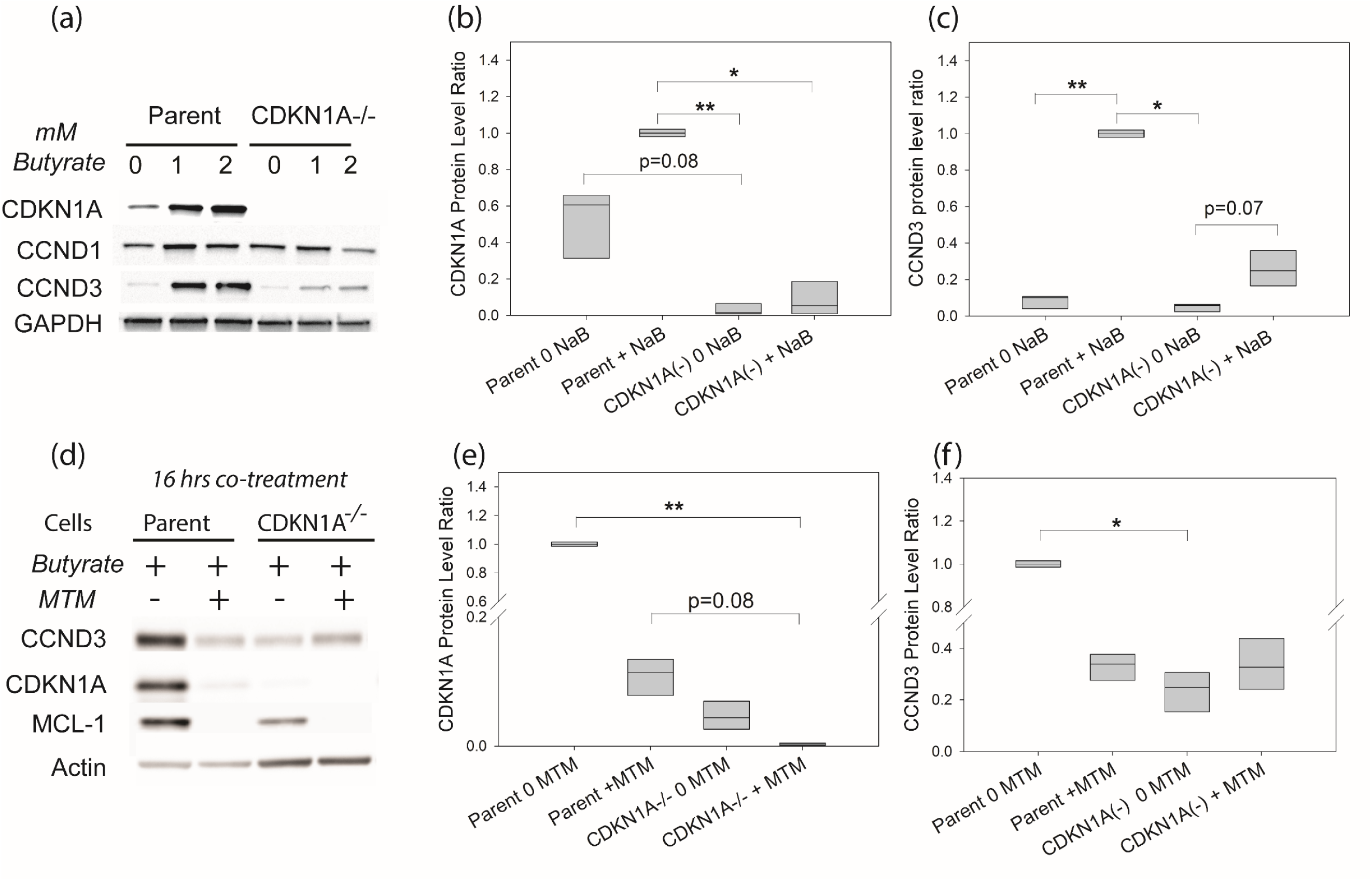
CCND3 protein level was dramatically reduced in butyrate-treated cells genetically or pharmacologically depleted in CDKN1A protein. (a)(b)(c) Parent and CDKN1A^-/-^ HCT116 cells were cultured in media containing initially different doses of butyrate during 16 hours. Extracted total proteins were separated as in Fig.1. (a) Images of a representative western analysis for CCND3, CCND1 and CDKN1A proteins. We show graphical representation of quantitative measurements of CCND3 (b) and CDKN1A (c) signals issued from three western analysis for [Butyrate]init = 2 mM. (c). A Kruskal-Wallis rank test indicated a significant difference between Cells + treatments for both CDKN1A(p=0.020) and CCND3 (P=0.019) proteins. Post hoc analyses with Holm correction showed that this effect was mainly due to differences between “Parent cells+2mM” butyrate with “CDKN1A^-/-^ cells + 0mM” butyrate (p =0.035) and with “CDKN1A^-/-^ cells+2mM” butyrate (p=0.079) for CDKN1A protein, and between “Parent cells+2mM” with “CDKN1A^-/-^ cells+0mM” butyrate (p=0.026) for CCND3. (d)(e)(f) Cells were cultured in a medium containing initially 1.5 mM butyrate in the presence of MTM or PBS. Extracted proteins were separated as in Fig.2. (d) Images of a representative western analysis. Graphical representation of quantitative measurements of CCND3 (b) and CDKN1A (c) signals issued from three different extracts for control or MTM-treated parent and CDKN1A^-/-^ cells analysis are shown. A Kruskal-Allis rank test indicated that the difference for CCND3 and CDKN1A protein level ratio between control and MTM-treated parental and CDKN1A-/-cells was significant (P = 0.016 and P = 0.003 respectively). Post hoc analyses with Holm correction showed that the MTM effect was mainly due to differences between “Parent cells+0MTM” with “CDKN1A^-/-^ cells + 500MTM” (p =0.002), while the difference between “Parent cells+0MTM” and “CDKN1A^-/-^ cells+500MTM” (p=0.081) for CDKN1A protein, and between “Parent cells+0MTM ” and “CDKN1A^-/-^ cells+0MTM” (p=0.010) for CCND3 were poorly significant.

To further confirm CDKN1A’s importance in CCND3 stabilization, we employed the antibiotic Mithramycin A (MTM), a known inhibitor of HDACi-induced CDKN1A transcription [73]. We reasoned that MTM treatment of parental cells would mimic the CDKN1A^-/-^ condition, allowing us to distinguish CDKN1A-dependent from CDKN1A-independent effects of MTM on CCND3.

In parental cells, MTM strongly reduced butyrate-induced CDKN1A expression, with a corresponding partial reduction in CCND3 protein levels (Fig. 3(d),(e)). In contrast, MTM treatment of CDKN1A^-/-^ cells did not further decrease CCND3; instead, it slightly but significantly (**P=0.032**) increased CCND3 levels (Fig. 3(d)). This stimulatory effect was specific since MTM still effectively reduced protein levels of another known MTM-sensitive target, Mcl-1, in both cell types [74]. Thus, the reduction of CCND3 in MTM-treated parental cells likely primarily resulted from reduced CDKN1A expression, while MTM exhibited a CDKN1A-independent stimulatory effect on CCND3 protein level.

Together, these results strongly support a model in which CDKN1A stabilizes CCND3 by protecting it from degradation, particularly during butyrate-induced cell-cycle arrest.

### 3.4. Impact of ectopic CCND3-HA expression on cellular responses to butyrate

To further assess the functional significance of increased CCND3 abundance during butyrate-induced cell-cycle arrest, we generated three HCT116 cell clones (L3, P3, and Q3) expressing varying levels of recombinant CCND3 tagged with hemagglutinin (HA), termed CCND3-HA (protein B in Fig. S1, Supplementary Data). Strikingly, increasing expression of recombinant CCND3-HA corresponded to decreased relative abundance of endogenous CCND3 in butyrate-treated cells (Fig. 4(a)), suggesting possible competition or feedback between recombinant and endogenous proteins.

**Fig. 4:**
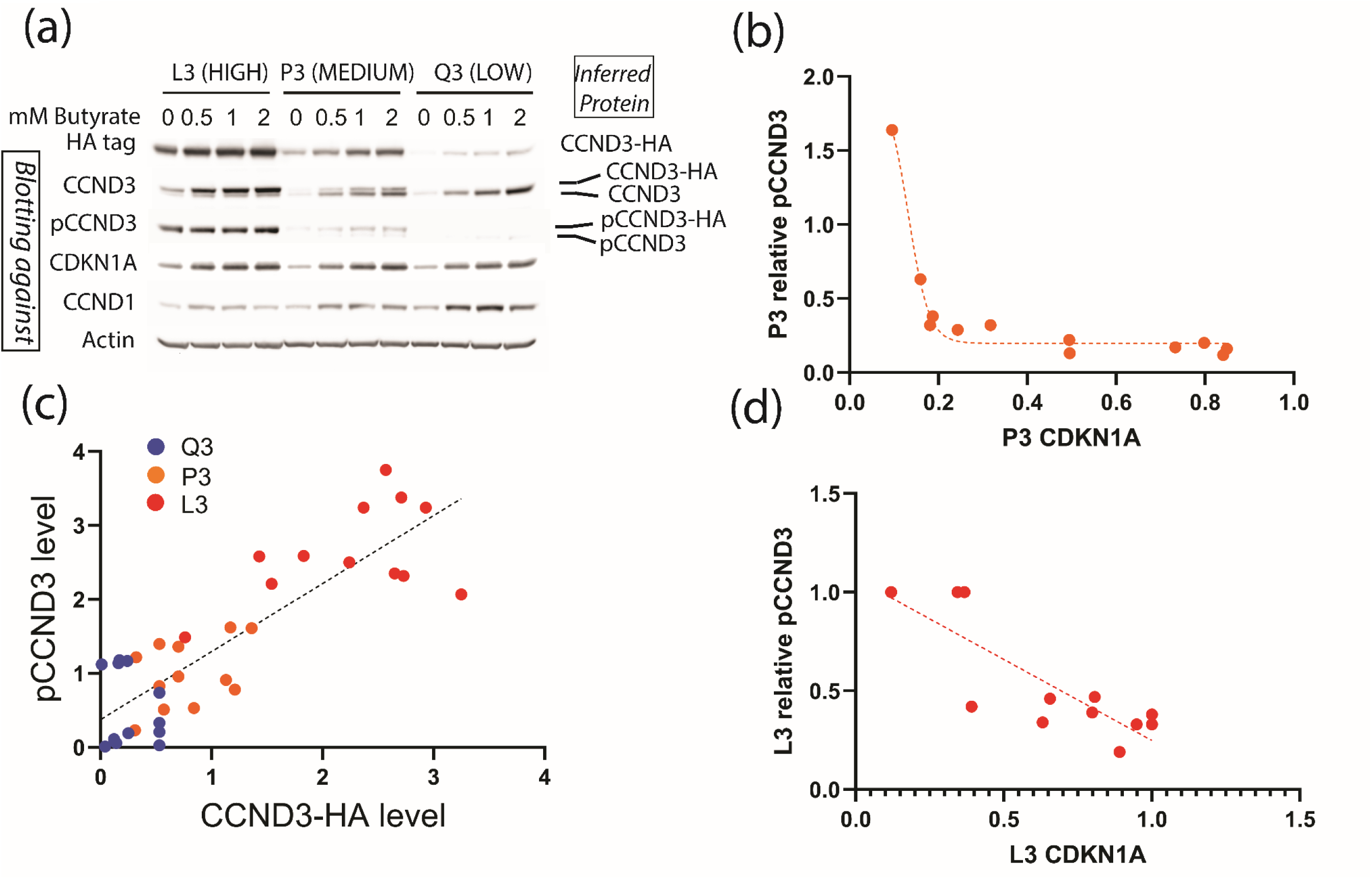
Fate of recombinant CCND3 tagged with Hemagglutinin HA sequence (CCND3-HA) that were expressed at different levels in HCT116 cells. The response to butyrate treatment in three clones, Q3 (LOW), P3 (MEDIUM) and L3 (HIGH) expressing LOW, MEDIUM and HIGH level of CCND3-HA respectively, was studied. Proteins were extracted as described in Fig.1. (a) Images of a representative western analysis. Results from 3 similar western analyses were used to generate graphics. They indicated (b) a significant correlation (Pearson coefficient r=0.842; P<0.001) between total phosphoCCND3 (native + recombinant) and CCND3-HA protein levels. In P3 cells (c) and L3 (d) cells, the relative phosphoCCND3 protein level (ratio=total phosphoCCND3/total (native + recombinant) CCND3 protein level) was inversely proportional to CDKN1A protein levels. A sigmoidal dose response and a linear decrease described this relationship in P3 (R^2^=0.979 but without complete CI) and L3 (R^2^= 0.677; P=0.001) cells respectively.

In the absence of butyrate, the growth rates of these clones were comparable to the parental HCT116 cell line at all fetal calf serum (FCS) concentrations tested (2.5%, 5%, and 10%; data not shown). After 5–6 days of exposure to 1 mM butyrate, all three clones showed reduced residual cell growth compared to parental cells but similar to each other (Fig. 5(a)). Specifically, at 2.5% FCS with 1 mM butyrate, residual growth ranged from 21% to 34% (Q3), 15% to 31% (P3), and 7% to 26% (L3). These differences did not reach statistical significance.

**Fig. 5:**
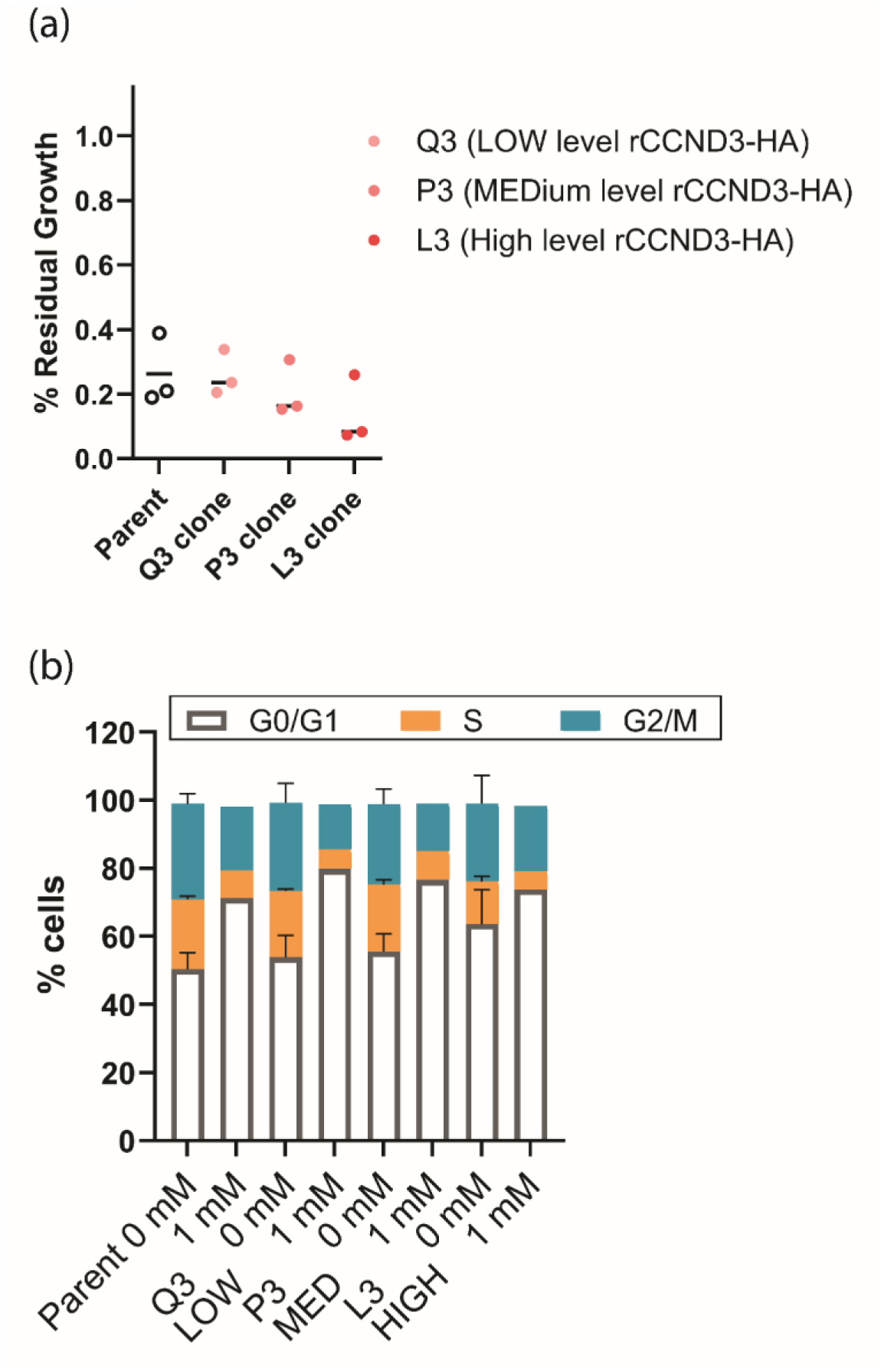
Consequences of ectopic expression of the recombinant CCND3-HA on butyrate-induced growth inhibition in HCT116 cells and on the cell cycle. All cells were cultured with 2.5% FCS. (A) Residual growth of HCT116 parental and clone cells. The percentage of remaining growth of parental and clone cells containing either LOW (Q3) MEDIUM, (P3) or HIGH (L3) level of recombinant CCND3-HA exposed to 1 mM butyrate was compared to the growth in control condition, without butyrate, of each cell population. The lines indicate the means of the three plotted values of maximal residual growth. (b): Cell phases of HCT116 parental and clone cells after 16-18 hours in 0 or 1 mM butyrate. The DNA content of individual 7-AAD-stained cells were determined with flow cytometry.

Consistent with these observations, cell cycle analyses performed under identical conditions (0 or 1 mM butyrate; Fig. 5(b)) revealed similar trends among clones. In untreated clones, the percentages of cells in the G1 phase were approximately 49% (Q3), 52% (P3), and 56% (L3), and those in S-phase were about 20% (Q3), 21% (P3), and 14% (L3). Following butyrate treatment, the percentage of cells in S-phase markedly decreased to averages of 6% (Q3), 8% (P3), and 5.5% (L3), with corresponding proportional increases in the G1 and G2 phases.

Although differences among clones were not statistically significant, clone L3 displayed a trend toward slightly reduced growth and fewer cells in S-phase. Given the known heterogeneity of the HCT116 cell line, these differences might however reflect genetic or phenotypic variations acquired during transfection or clonal selection.

Increased recombinant CCND3-HA levels had therefore only modest effects on proliferation in absence of butyrate and on cell-cycle arrest induced by butyrate. These results indicated that elevated cyclin D3 expression in HCT116 cells did not significantly contribute to inhibit proliferation, contrary to what has been observed in a few cases [75, 76].

Further analyses of CCND3 phosphorylation revealed additional insights. In clone Q3, expressing the lowest recombinant CCND3-HA level, phosphorylated CCND3 (endogenous or recombinant) was scarcely detected, similar to parental cells (Fig. 1(c), Fig. 4(a)). Conversely, in clone L3, abundant CCND3-HA allowed clear detection of its phosphorylated form. Intriguingly, despite endogenous CCND3 being predominant in clone P3, phosphorylation occurred preferentially on the recombinant CCND3-HA. Quantitative analyses revealed a strong positive correlation (Pearson coefficient r = 0.842; P<0.001) between total phosphorylated CCND3 (native and recombinant) and recombinant CCND3-HA levels (Fig. 4(b)). Additionally, the relative phosphorylation of total CCND3 proteins inversely correlated with CDKN1A levels in clones L3 and P3 (Fig. 4(c),(d)). Specifically, phosphorylation was highest in untreated cells where CDKN1A was lowest, suggesting elevated CDKN1A may partially inhibit CCND3 phosphorylation. Notably, CDKN1A expression profiles remained consistent across clones (Fig. 4(a)), indicating CCND3 levels did not reciprocally affect CDKN1A abundance.

Immunofluorescence analysis further indicated that endogenous CCND3 localized primarily to the nucleus of parental HCT116 cells after 16 hours of butyrate treatment (Fig. 6(a)). In untreated asynchronous cells, CCND3 signals were weak and distributed in both nuclear and cytoplasmic compartments. After butyrate exposure, nuclear accumulation of CCND3 markedly increased. Similarly, recombinant CCND3-HA predominantly localized to nuclei in butyrate-treated clones, as demonstrated by anti-HA antibody staining (Fig. 6(b)). Moreover, nuclear localization correlated positively with recombinant CCND3-HA abundance across clones (L3 > P3 > Q3), confirming substantial nuclear retention of recombinant CCND3 following butyrate treatment.

**Fig. 6:**
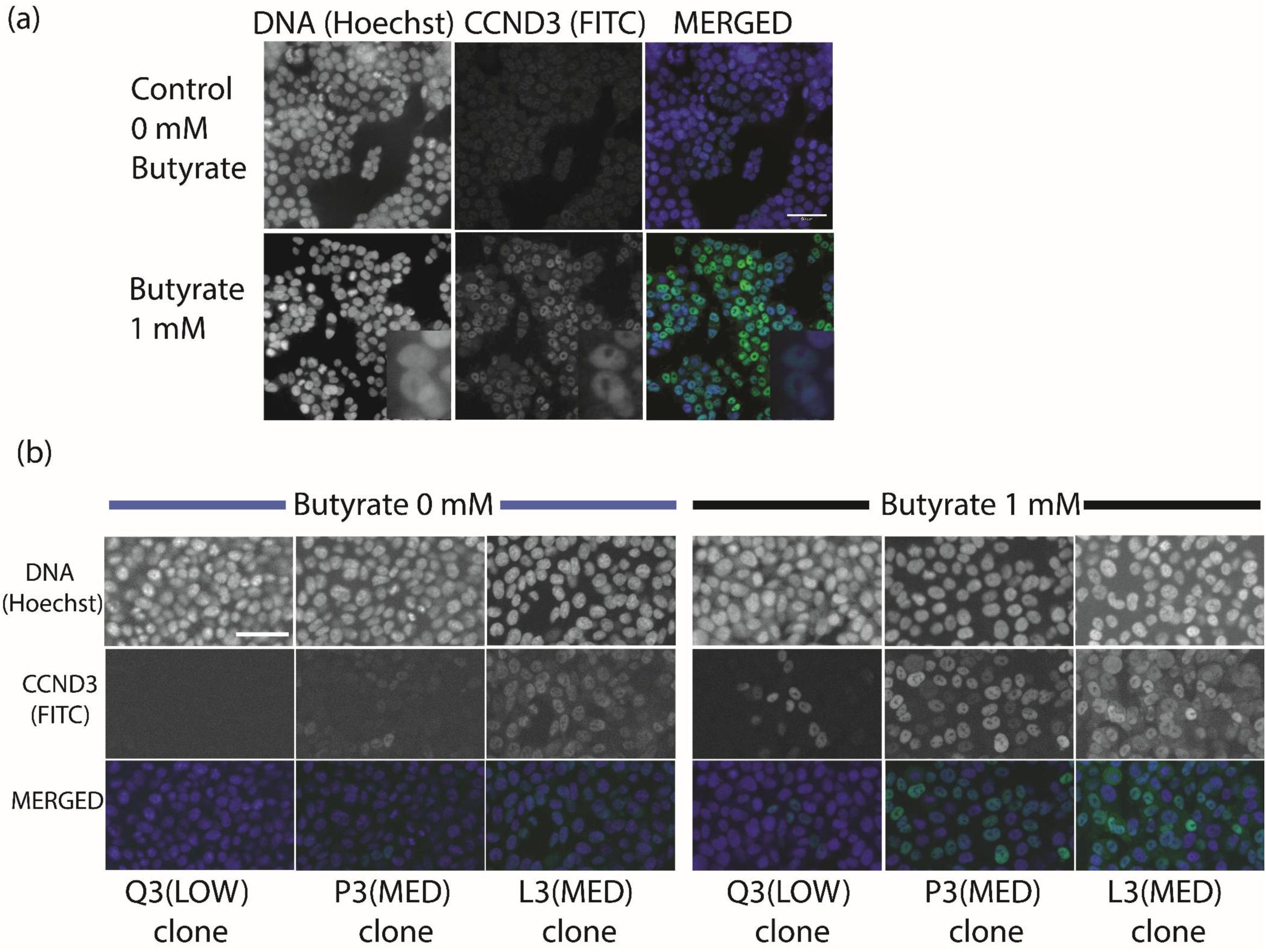
Immunofluorescence detection of CCND3 and CCND3-HA protein in HCT116 (a) parental and (b) clones cells expressing a transgene coding for CCND3-HA and expressed at different levels as detailed in Fig.6. HCT116 parental (a) and HCT116 Q3, P3 and L3 clone(b) cells were exposed to 0 and 1 mM butyrate for 24 hours. After the treatments, cells cultured in 96-well plates were fixed and probed with anti-CCND3 antibodies and FITC-labeled secondary antibodies. HOECHST: DNA counter-staining with Hoechst 33342; FITC: immune-labeling with primary CCND3 and FITC-coupled secondary antibodies. Merged: merged images of blue (HOECHST) and green (FITC) signals. In (a) enlarged views of the specified region are shown in inserts. Scale bar stands for 50µm. Observations were performed on HCS microscope in confocal mode with the x20 objective.

Collectively, these results indicate that ectopic CCND3-HA expression minimally impacts cell proliferation or cell-cycle arrest induced by butyrate. This limited impact likely arises from either the absence of reciprocal feedback regulation of CDKN1A or partial functional differences between recombinant and endogenous CCND3 proteins, as discussed further below.

### 3.5. CDKN1A physically interacts with CCND3 complexes in butyrate-treated cells

Given the strong influence of CDKN1A levels on CCND3 stability, we examined whether CDKN1A physically associates with CCND3 in butyrate-treated cells using co-immunoprecipitation (co-IP) methods.

Using anti-HA antibodies in clones expressing recombinant CCND3-HA (L3 and P3), we found that CDKN1A co-immunoprecipitated specifically with CCND3-HA (Fig. 7(a)). No precipitation of either CCND3 or CDKN1A was observed in parental cells, confirming specificity. The amounts of co-immunoprecipitated CCND3 and CDKN1A correlated proportionally with their abundance in the input extracts, both in the presence and absence of butyrate. However, the ratio of co-immunoprecipitated CCND3 to CDKN1A differed from their ratios in input extracts, suggesting that only a fraction of total CDKN1A physically interacts with CCND3.

**Fig. 7:**
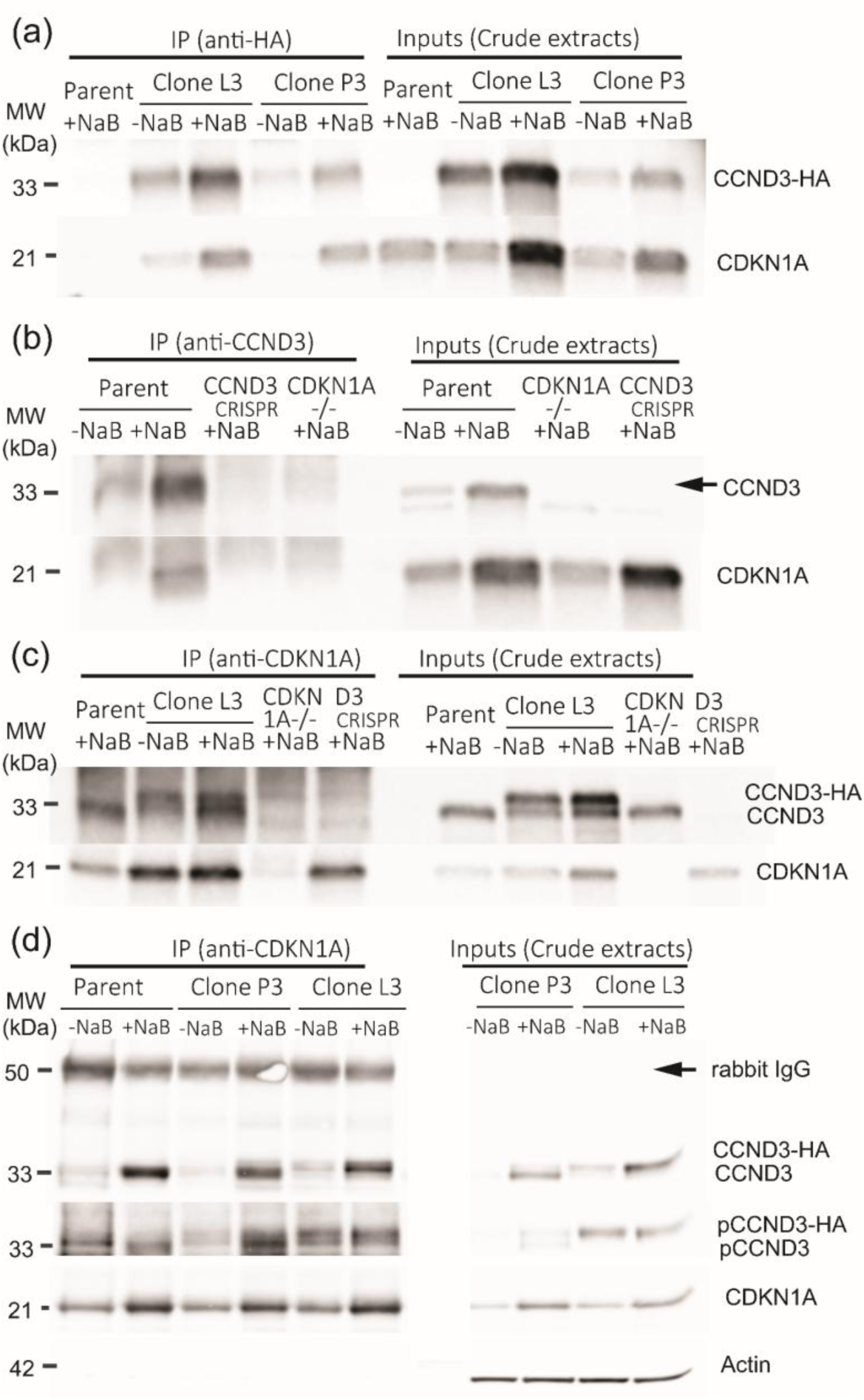
CCND3 and CDKN1A co-immunoprecipitated (co-IP) in butyrate-treated HCT116 cells. Co-immunoprecipitation targeted (a) CCND3-HA with anti-HA antibodies, (b) CCND3 with anti-CCND3 antibodies, and (c) (d) CDKN1A with anti-CDKN1A antibodies. Immunoprecipated (IP) fractions were analyzed on western blots and compared to the “crude” protein extracts used for co-IP. Each type of IP was performed at least twice.

We further confirmed these interactions using antibodies directed against the native CCND3 C-terminal region in parental cells. Both CCND3 and CDKN1A proteins co-immunoprecipitated successfully from parental cell extracts (Fig. 7(b)). Importantly, CDKN1A did not co-immunoprecipitate from extracts of a CRISPR/Cas9-generated mutant lacking CCND3’s C-terminal region (Fig. 7(b)) highlighting the essential nature of this region for CDKN1A binding.

To validate the specificity of the CCND3–CDKN1A interaction further, we conducted reciprocal immunoprecipitation using anti-CDKN1A antibodies. Neither CCND3 nor CDKN1A were recovered from CDKN1A^-/-^ cell extracts, confirming the immunoprecipitation specificity (Fig. 7(c),(d)). In contrast, endogenous CCND3 co-immunoprecipitated with CDKN1A in parental cells, and both endogenous CCND3 and recombinant CCND3-HA were co-immunoprecipitated with CDKN1A from clone L3 and P3 cell extracts. Interestingly, a fraction of co-immunoprecipitated CCND3 (endogenous and recombinant) was phosphorylated at Thr283 (Fig. 7(d)). However, our attempts to quantify precise phosphorylation ratios using LC-MS have not yet succeeded.

Collectively, these findings strongly support a model in which CCND3 and a subset of CDKN1A form stable physical complexes in the nuclei of butyrate-treated HCT116 cells, which are largely arrested in the cell cycle. Previous studies have demonstrated that CDKN1A physically interacts with cyclin D proteins, promoting the formation, nuclear import, and activity of cyclin D–CDK4/6 complexes. Considering these observations and the extensive intrinsically disordered regions (IDRs) present in CDKN1A—and smaller IDRs in CCND3— we next sought to computationally model their structural interaction based on available partial structural data.

### 3.6. Proteomic identification of CCND3-associated proteins reveals potential new interactors, including CDK5

To further elucidate CCND3’s functional interactions in butyrate-treated cells, we identified additional proteins co-immunoprecipitating with recombinant CCND3-HA using LC-MS analysis. Samples from L3 and P3 clones cultured with or without butyrate treatment—each analyzed in triplicate—were subjected to rigorous selection criteria to ensure robust identification (Materials and Methods and Supplementary Data). We verified that identified proteins were not present in parental control samples or detectable as alternative isoforms. This stringent approach identified three proteins consistently associated with CCND3-HA in untreated cells: CDKN1A (confirming earlier western blot data), and the known CCND interactors CDK4 and CDK6 (Table 1). Comparing samples expressing similar recombinant CCND3-HA levels (untreated L3 versus butyrate-treated P3 cells) revealed that four proteins (CDKN1A, CDK4, CDK6, and CDK5) were more abundant in butyrate-treated samples. Notably, CDK5 was uniquely detected in co-immunoprecipitates from butyrate-treated cells (Table 1), suggesting a possible butyrate-specific interaction.

**Table 1:** List of proteins co-immunoprecipitated with the recombinant protein CCND3-HA in L3 cells without NaB and P3 cells exposed to NaB. Both cell types expressed similar levels of the recombinant protein.

| Proteins interacting with CCND3-HA | Swiss-Prot record | Without butyrate (PAI L3 cells) | With butyrate (PAI P3 cells) | Description and Gene Name (GN) | Subcellular localization |
| --- | --- | --- | --- | --- | --- |
| CDKN1A | P38936\|CDN1A_HUMAN | - (1.33±0.22) | + (2.70±0.04) | Cyclin-dependent kinase inhibitor 1 OS=Homo sapiens GN=CDKN1A | <u>Nucleus</u>; cytoplasm |
| CDK4 | P11802\|CDK4_HUMAN | - (0.90±0.22) | + (1.26±0.04) | Cyclin-dependent kinase 4 OS=Homo sapiens GN=CDK4 | <u>Nucleus</u>; cytoplasm |
| CDK5 | Q00535\|CDK5_HUMAN | - (0.06±0.11) | + (0.75±0.04) | Cyclin-dependent-like kinase 5 OS=Homo sapiens GN=CDK5 | <u>Nucleus</u>; cytoplasm |
| CDK6 | Q00534\|CDK6_HUMAN | - (0.09±0.07) | + (0.58±0.04) | Cyclin-dependent kinase 6 OS=Homo sapiens GN=CDK6 | <u>Nucleus</u>; cytoplasm |
| CCND3-HA |  | + (0.64±0.07) | +(0.62±0.04) | Cyclin D3 with a C-terminal HA tag | nucleus |

CDK5 is best characterized for its high expression and functional roles in the adult mouse brain, where it regulates diverse physiological processes in post-mitotic neurons. However, CDK5 is also expressed in various other cell types, phosphorylating substrates such as the retinoblastoma protein (Rb) (reviewed by [77]. Although CDK5 activation typically depends on its association with the non-cyclin activators CDK5R1 (p35) and CDK5R2 (p39), its activity can also be inhibited by cyclins including CCND1 and the CDKN1B–CCNE complex. Dysregulated CDK5 activity has been implicated in abnormal neuronal cell-cycle reentry and apoptosis in Alzheimer’s disease, and CDK5 overexpression is frequently observed in numerous cancers. Moreover, CDK5 phosphorylates the guanine nucleotide exchange factor (GEF) GIV, modulating the balance between cell proliferation and migration [78].

In colorectal cancer specifically, a functional role for CDK5 in promoting cell proliferation and invasion has been established [79]. Knockdown of CDK5 significantly reduced both proliferation and invasion in HCT116 cells [79]. This inhibitory effect primarily resulted from decreased ERK5-AP-1 signaling, an emerging hallmark pathway in cancer progression [80, 81], since ERK5 activation depends on phosphorylation at Thr732 by CDK5 [79].

Thus, identifying CDK5 among CCND3-associated proteins, specifically enriched in response to butyrate treatment, provides novel insights into potential molecular interactions contributing to butyrate-induced cell-cycle arrest in colorectal cancer cells.

### 3.7. Structural modeling predicts a mechanistic basis for CDKN1A-mediated stabilization of CCND3

To gain mechanistic insights into how CDKN1A stabilizes CCND3, we performed computational modeling of the CDKN1A–CCND–CDK4 complexes using existing structural data combined with AlphaFold2 (AF2) predictions. Although the CDKN1A–CCND1–CDK4 complex structure (PDB ID: 6P8H) was previously resolved, critical regions—particularly intrinsically disordered regions (IDRs) of CDKN1A and the terminal regions of CCND1— remained unresolved (Fig. S12 (a) and (b), Supplementary Data). These unresolved regions include the functionally important RRL motif in CDKN1A and the conserved C-terminal Thr residues in cyclins (Thr286 in CCND1, Thr283 in CCND3), both implicated in phosphorylation-dependent nuclear export and degradation.

Leveraging AF2 structural predictions, we generated complete atomic models for CDKN1A– CCND1–CDK4 and CDKN1A–CCND3–CDK4 complexes, including previously unresolved IDRs. AF2-predicted structures showed excellent alignment with existing crystallographic data (backbone RMSDs: CDK4, 0.692 Å; CDKN1A, 0.521 Å; CCND1, 0.472 Å; CCND3, 0.639 Å) (Fig. 8). Surprisingly, both Thr286 (CCND1) and Thr283 (CCND3) were predicted to localize near CDKN1A’s RRL motif. However, the predicted structural conformations of the cyclins’ C-terminal tails differed substantially: AF2 predicted two short helices within CCND1’s tail, whereas CCND3’s tail lacked clearly defined secondary structures. Both cyclins contained (ANCHOR2 profiles in Fig. S12 (b) and (c), Supplementary Data) predicted molecular recognition features (MoRFs) around these critical Thr residues (CCND1 residues 278–295, CCND3 residues 281–292), with CCND3 containing an additional MoRF (residues 249–259).

**Fig. 8:**
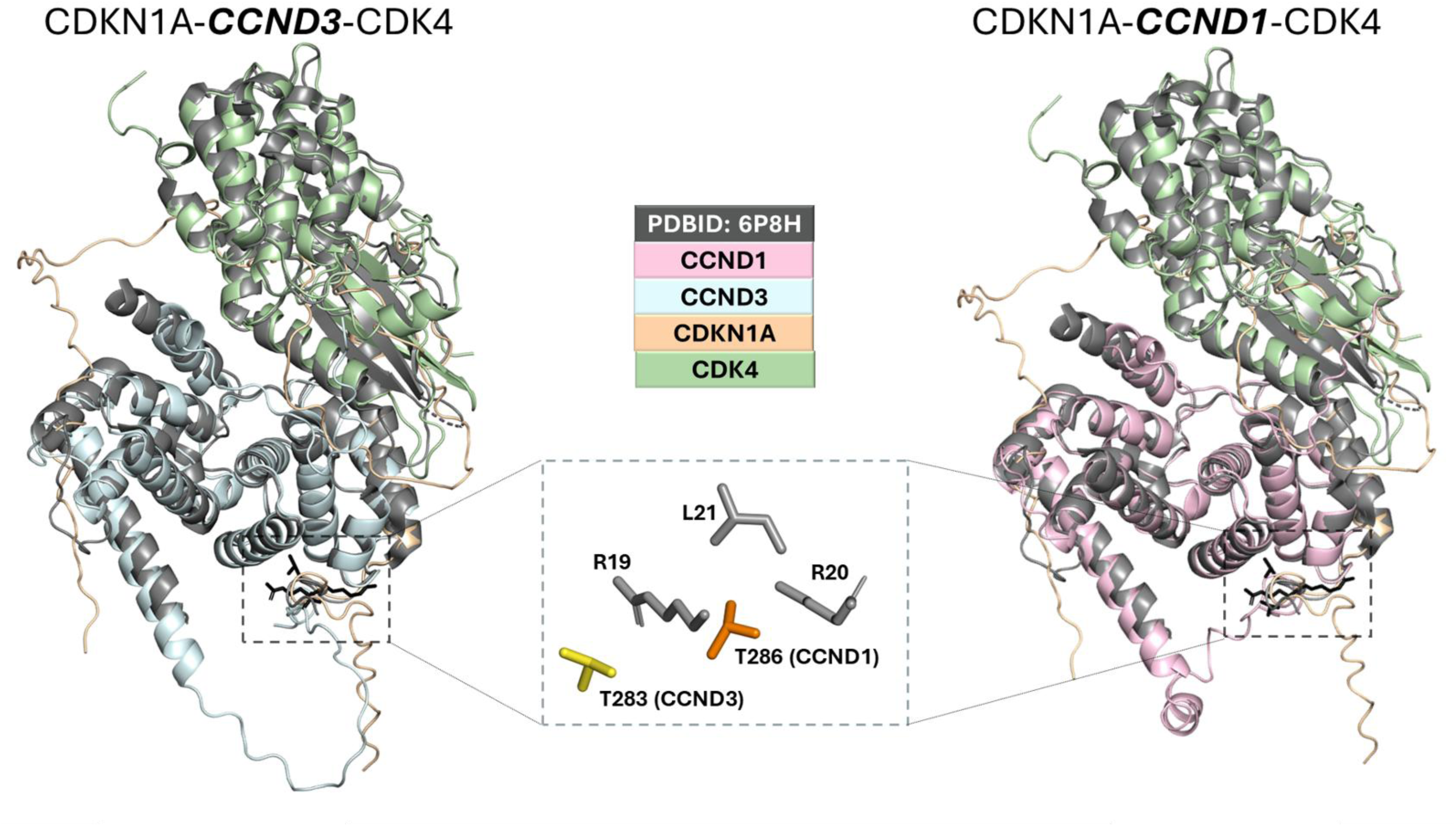
Structural comparison of the crystallographic (PDB ID: 6P8H) and AlphaFold2-predicted (AF2) CDKN1A–CCND1/3–CDK4 complexes. Each subunit (CDKN1A, CCND1/CCND3, and CDK4) was predicted separately by AF2 and then overlaid onto the experimentally determined partial complex. Shown are the reassembled AF2-predicted CDKN1A–CCND3–CDK4 (left) and CDKN1A–CCND1–CDK4 (right) structures, highlighting how unresolved intrinsically disordered regions (IDRs) and extended C-terminal tails differ from the crystal structure. Dashed boxes mark the spatial proximity between CDKN1A’s RRL motif (Arg19–Arg20–Leu21) and the conserved cyclin Thr residues (Thr283 in CCND3 or Thr286 in CCND1). The inset magnifies these residues (RRL in gray; Thr283 in red; Thr286 in yellow), showing side-chain orientations. Notably, AF2 predicts variation in the C-terminal tails, suggesting CCND1 and CCND3 may exhibit distinct molecular recognition features (MoRFs).

To resolve ambiguities arising from these structural differences, we conducted molecular dynamics (MD) simulations of CDKN1A–CCND1–CDK4 and CDKN1A–CCND3–CDK4 complexes. Simulations (300 ns) showed stable interactions consistent with experimental data. In both complexes, ordered regions of CDK4 remained structurally stable, with minimal variation except at its C-terminal tail. Notably, the CDK4 tail was stabilized by interactions with CDKN1A residues in the CCND1 complex but remained flexible and solvent-exposed in the CCND3 complex.

CDKN1A adopted distinct conformations depending on the cyclin partner. In the CCND1 complex, CDKN1A exhibited minimal secondary structure, whereas, in the CCND3 complex, extended helices formed. Furthermore, although the RRL motif of CDKN1A was located near the conserved Thr residues of both cyclins, direct hydrogen bonding was observed exclusively with CCND1. In contrast, no direct interactions occurred between the RRL motif and CCND3-Thr283. MD analyses revealed a striking difference in solvent accessibility of the Thr residues: CCND1-Thr286 was partially solvent-exposed (mean solvent-accessible surface area, SASA = 0.61 ± 0.16 Å²), whereas CCND3-Thr283 was buried (SASA = 0.09 ± 0.06 Å²) (Fig. 9). This suggests CDKN1A binding induces structural rearrangements in CCND3, burying Thr283 and thereby preventing its phosphorylation and subsequent nuclear export. Experimentally, we observed that both native and recombinant CCND3 co-immunoprecipitated with CDKN1A existed as phosphorylated and unphosphorylated forms, though the unphosphorylated form predominated (Fig. 1(c), Fig. 7(d)). Two scenarios could explain this observation: (1) phosphorylation does not disrupt CCND3–CDKN1A interactions significantly; or (2) CDKN1A binding actively inhibits Thr283 phosphorylation, but a minor subpopulation escapes inhibition. The behavior of recombinant CCND3-HA supports the second scenario. Increased CCND3-HA expression correlated strongly (r = 0.842; P<0.001) with increased phosphorylation (Fig. 4(b)), reaching maximal levels when CDKN1A abundance was lowest (Fig. 4(c),(d)). Moreover, phosphorylated CCND3-HA levels in complex with CDKN1A remained constant irrespective of butyrate treatment, suggesting preferential CDKN1A association with a minor phosphorylated subpopulation (Fig. 7(d)). Collectively, these findings strongly indicate CDKN1A binding inhibits Thr283 phosphorylation.

**Fig. 9.**
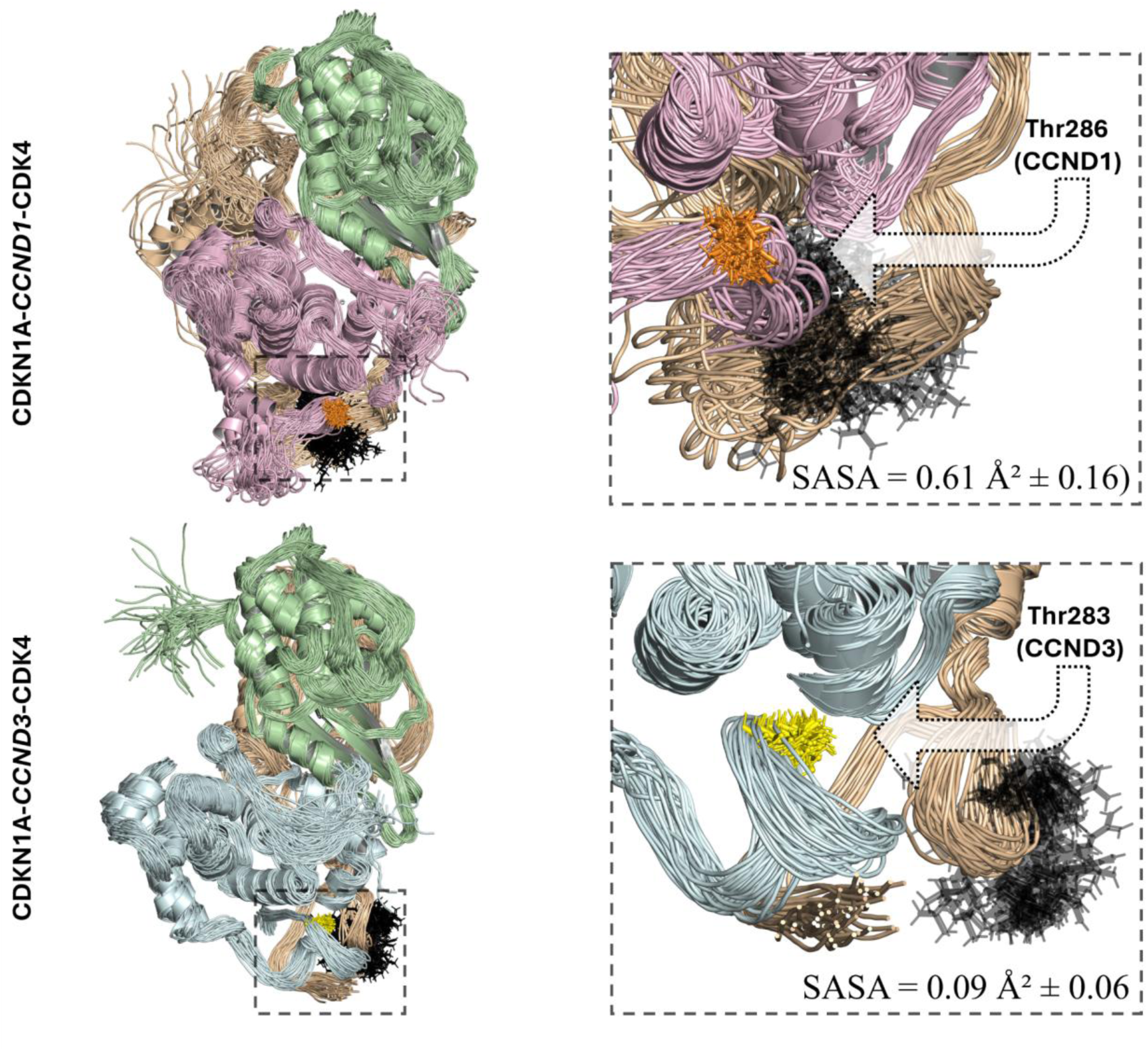
Molecular dynamics (MD) ensembles of the final 100 ns of the CDKN1A–CCND1–CDK4 (top) and CDKN1A–CCND3–CDK4 (bottom) complexes. Each ensemble comprises 50 snapshots taken at 2 ns intervals, illustrating how CDKN1A (wheat) interacts distinctly with CCND1 (pink) versus CCND3 (blue); CDK4 is shown in green. A dashed rectangle highlights the region containing the RRL motif of CDKN1A (black) and the conserved cyclin Thr residues (Thr286 in CCND1, orange; Thr283 in CCND3, yellow), whose side chains are displayed. The right panels show zoomed-in views of these residues, with calculated solvent-accessible surface areas (SASA) indicating Thr286 is partially solvent-exposed (0.61 ± 0.16 Å^2^), whereas Thr283 is nearly buried (0.09 ± 0.06 Å^2^). In the CDKN1A–CCND1 complex, the RRL motif forms multiple hydrogen bonds with CCND1, orienting Thr286 outward and facilitating phosphorylation. By contrast, Thr283 in CCND3 remains solvent-inaccessible, suggesting that CDKN1A binding induces a locally folded conformation that hinders phosphorylation and promotes CCND3 stabilization.

Thus, structural modeling combined with experimental data supports a mechanistic model where CDKN1A binding induces conformational changes in CCND3’s C-terminal tail, burying Thr283 and preventing its phosphorylation-dependent nuclear export and degradation. In contrast, distinct structural interactions in CCND1 may allow Thr286 phosphorylation, highlighting cyclin-specific regulatory roles of CDKN1A. Furthermore, the recombinant HA-tagged CCND3 might alter native conformations, partially disrupting normal CDKN1A interactions and enhancing Thr283 accessibility.

### 3.8. Proposed model of CDKN1A-mediated CCND3 stabilization and implications for cell-cycle arrest

Our combined experimental and computational data support a comprehensive model in which CDKN1A (p21) significantly stabilizes cyclin D3 (CCND3) protein following butyrate treatment, thereby contributing directly to cell-cycle arrest at the G1 phase. Previous studies demonstrated that CDKN1A could inhibit nuclear export and subsequent degradation of CCND1 [32], possibly favoring cell-cycle progression when CDKN1A levels remain low. Indeed, cyclin D–CDK4 complexes can sequester CDKN1A/B, preventing CDK2 inhibition and facilitating G1/S transition [20]. Our results extend these findings by identifying a cyclin-specific role for CDKN1A in regulating CCND3 stability during butyrate-induced cell-cycle arrest.

Specifically, our experimental findings showed that CDKN1A directly or indirectly associates with CCND3 in nuclear complexes enriched upon butyrate treatment (Fig. 7). Proteomic analysis identified additional CCND3-associated proteins, including CDK4, CDK6, and importantly, CDK5—a kinase linked to colorectal cancer cell proliferation and migration [78, 79]. The identification of CDK5 raises intriguing possibilities about novel regulatory interactions involving CCND3–CDKN1A complexes in cell-cycle arrest and differentiation pathways.

Complementary structural modeling and molecular dynamics simulations further revealed how CDKN1A binding to CCND3 could inhibit its phosphorylation-dependent nuclear export. Computational predictions indicated that CDKN1A binding triggers conformational rearrangements in CCND3’s intrinsically disordered C-terminal tail, burying the conserved Thr283 residue, thereby preventing efficient phosphorylation (Fig. 9). These results were strongly supported by experimental evidence showing preferential stabilization of unphosphorylated CCND3 in the presence of elevated CDKN1A levels (Fig. 1(c) and Fig. 4(b,d)).

Based on these observations, we propose a model in which elevated CDKN1A levels induced by butyrate stabilize CCND3 predominantly in an unphosphorylated, nuclear-retained form. This mechanism would thus contribute directly to butyrate-mediated cell-cycle arrest in colorectal cancer cells by preventing CCND3 degradation and maintaining inhibitory cyclin D–CDK–CDKN1A complexes within the nucleus.

Furthermore, considering the intrinsically disordered nature of CDKN1A and its ability to drive liquid-liquid phase separation (pLLPS ≥ 0.60) [82–85], our findings also open the possibility that phase-separated condensates formed by CDKN1A could regulate cyclin-CDK complex localization, stability, and activity. Such biomolecular condensates might thus play critical but currently unexplored roles in the cellular mechanisms underlying butyrate-induced arrest in cancer and normal stem cells.

Overall, our findings provide novel mechanistic insights into the tumor-suppressive actions of butyrate through the CDKN1A-dependent stabilization of CCND3. Future investigations into the precise role of condensate formation and the functional significance of newly identified interactors such as CDK5 may yield further insights into colorectal cancer biology and therapy.

## 4. Conclusion

Our study demonstrates a previously unexplored role for CDKN1A (p21) in mediating the paradoxical stabilization of cyclin D3 (CCND3) by preventing its phosphorylation-dependent nuclear export and subsequent degradation in colorectal cancer cells treated with butyrate. Whether this stabilization contributes directly to the G1-phase cell-cycle arrest induced by butyrate or to another process remains to be determined, but it provides new insights into the preventive action of dietary fiber-derived butyrate against colorectal carcinogenesis.

We identified a novel protein interaction involving CCND3, namely the association with CDK5, suggesting that CDKN1A–CCND3 complexes may have broader functional significance beyond canonical cyclin–CDK pathways. Structural modeling and molecular dynamics simulations further supported a mechanistic model wherein CDKN1A binding induces structural rearrangements in CCND3’s intrinsically disordered regions, limiting phosphorylation and nuclear export of CCND3.

Our findings also raise intriguing possibilities about the biological relevance of biomolecular condensates, driven by CDKN1A’s intrinsically disordered nature, in regulating cyclin–CDK complex stability and activity. Future investigations should explore the potential involvement of liquid-liquid phase separation (LLPS) and condensate formation in cell-cycle regulation, differentiation, and tumor suppression pathways.

In conclusion, these results expand our understanding of CDKN1A as a multifunctional regulator of cyclin stability, identify previously unrecognized protein interactions relevant to colorectal cancer biology, and highlight important new avenues for therapeutic exploration targeting CCND3-CDKN1A complexes.

## Supplementary data

The supplementary data file Ghrib_Dayhoff_supplementary.pdf can be uploaded The mass spectrometry proteomics data have been deposited to the ProteomeXchange Consortium via the PRIDE (https://www.ebi.ac.uk/pride/) partner repository with the dataset identifier PXD062450 and will become freely available once the paper has been accepted. The R studio script that was used to analyze these data can be obtained on request.

## Supporting information

The supplementary data file Ghrib_Dayhoff_supplementary_data.pdf can be uploaded

## Acknowledgements

A.Y. is grateful to the French Embassy in Cairo, Arab Republic of Egypt. C.C thanks ENIS in Sfax, Republic of Tunisia, for allowing him to complete his final 6-month-internship in Jouy-en-Josas, France. We are very grateful to Dr Celia Chamignon (Novobiome, France) for teaching RT-qPCR methods to M.A.G, to Drs Charlene Lasgi (Curie Institute, Orsay, France) and Aurelie Magniez (INRAe) for their help and advices in cytometry analysis. We thank Drs Pierre-Marie Girard (Curie Institute, Orsay, France) and Sylvie Demignot (EPHE and St-Antoine Research Center, Paris, France) for helpful discussions and Dr Pierre-Marie Girard for critical reading of a first draft. We thank Dr. Sameer Varma (University of South Florida, Tampa, USA) for supporting G.S. during this work.

## Author Contributions

**Mohammed A Ghrib**: investigation. **Guy W Dayhoff**: investigation, conceptualization, writing-review and editing. **Cherif Chahtour**: investigation, visualization. **Carine Rodrigues-Machado**: investigation, data curation, formal analysis. **Abdelrahaman Youssef**: investigation. **Giorgio Schillaci**: investigation. **Guillaume Gautreau**: formal analysis, visualization. **Céline Henry**: methodology. **Vladimir N. Uversky**: Investigation, conceptualization, writing-review and editing. **Pascal Rigolet**: formal analysis, visualization, writing-review and editing. **Hervé M. Blottiere**: investigation, conceptualization. **Jean-Marc Lelièvre**: investigation, conceptualization, formal analysis, visualization, supervision, writing-original draft.

## Conflicts of interest

The authors declare nο competing interests.

### Abbreviations

AF2: AlphaFold 2;
CCND1/2/3: Cyclin D1 D2/D3;
CDK1/2/4/5/6: cyclin-dependent kinases 1/2/4/5/6;
CDKN1A/B/C: CDK inhibitor1A/B/C also known as p21^Cip/Waf1^, p27^Kip1^ and p57^Kip2^ respectively;
CI: confidence interval;
DSB: double-strand break;
EC50: Half maximal effective concentration;
H/KDACi: Histone/Lysine deacetylase inhibitor;
IC50: Half maximal inhibitory concentration;
IDR/P: Intrinsically disordered region/protein;
LC-MS: liquid chromatography coupled to mass spectrometry;
LLPS: liquid-liquid phase separation (pLLPS= LLPS propensity score);
MoRF: Molecular recognition features;
NaB: Sodium butyrate;
NSAF: Normalized Spectral Abundance Factor (LC-MS data).
PAI: Protein Abundance Index (LC-MS data).

