## Supplementary material for "CDKN1A (p21^Cip/Waf1^) stabilizes Cyclin D3 by inhibiting its phosphorylation-dependent nuclear export following butyrate treatment": The supplementary data file Ghrib_Dayhoff_supplementary_data.pdf can be uploaded

**Ghrib, Dayhoff et al.. 2025**

**Fig. S1: CCND3 protein constructs used in this study**: (A) the endogenous human CCND3 protein, (B) the recombinant CCND3-HA protein and (C)(the putative truncated CCND3 proteins encoded after CRISPR mutagenesis. Rb refers to the N-ter domain that contains the binding region to Retinoblastoma protein; the CDK domain refers to the region that binds CDKs; the Repressor domain refers to the domain that was reported to bind p300 Histone/Lysine acetyl transferase H/KAT protein associated with gene transcription; HA refers to the Hemaglutinin HA tag. The arrow shows the peptide site corresponding to the RNA guide successfully used for CRISPR-Cas9-mediated mutagenesis of CCND3 genes. Key amino acid residues are indicated. In red boxes and letters indicates the epitope regions and the corresponding antibodies used in this study. C corresponds to a putative protein following partial repair of the CAS9 targeted site that changed the ORF of the 3’end of the CCND3 gene.

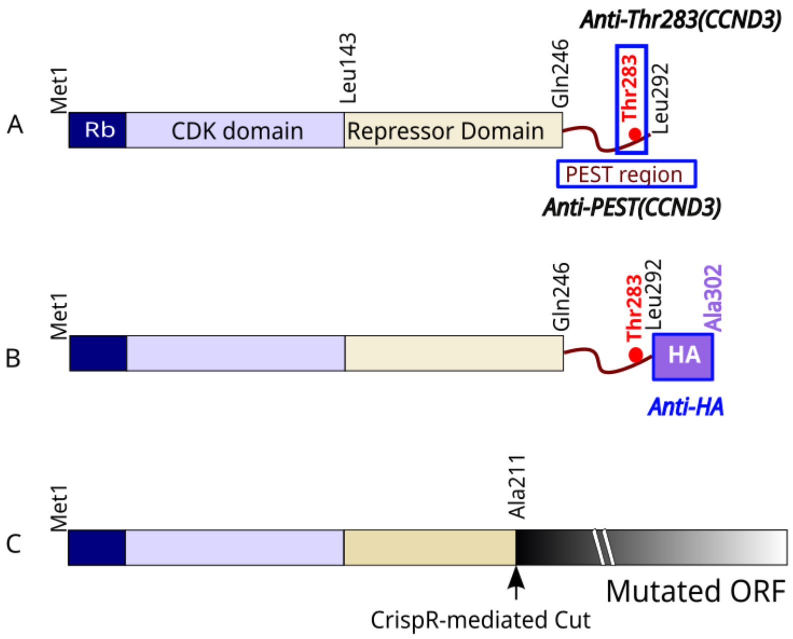

**Fig. S2**: **Western analysis of type D Cyclins  in Caco-2 cells**. Proteins were extracted as in Fig.1 from cells exposed to increasing concentrations of butyrate. Two blots of the same proteins extracts had to be run to avoid confusing signal between CCND2 and CCND3.

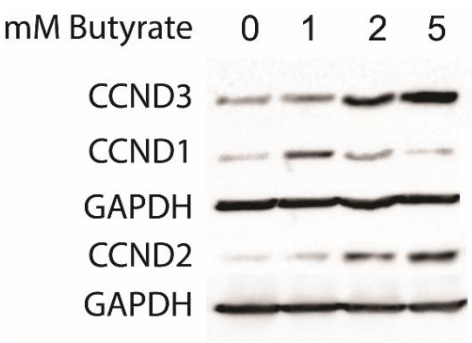

**Fig.S3**: **Graphical representation of the Ln values corresponding to western analyses shown in Fig.3(a)** for (CCND1) and(CCND3). The quantified signals allowed us to verify the linear fit of the decrease in protein level ratio vs Time for CCND1 (P= 0.0112) and CCND3 (P<0.0001) in butyrate-pretreated cells exposed to CHX, and to infer their half-life.

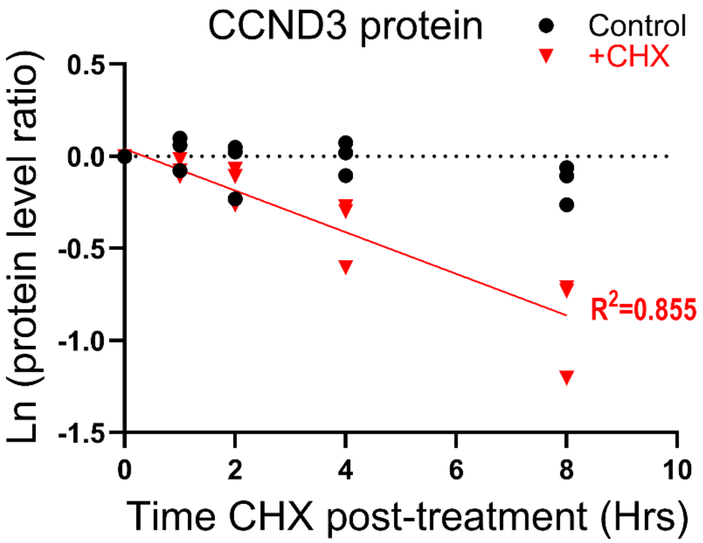


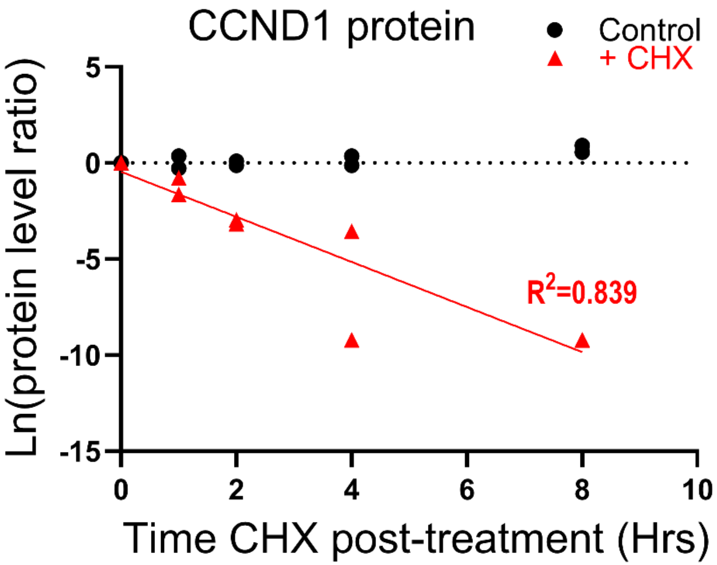

### Supplementary data related to the analysis of the proteomic data.

**Fig. S4**: **Screening protocol of proteins co-immunoprecipitated with anti-HA antibodies**. A rigorous screening algorithm was applied for this study, which led to retain proteins common to the six samples of L3/P3 transgenic cells cultured without NaB only, and proteins found in the six samples of L3/P3 cells cultured with NaB only.

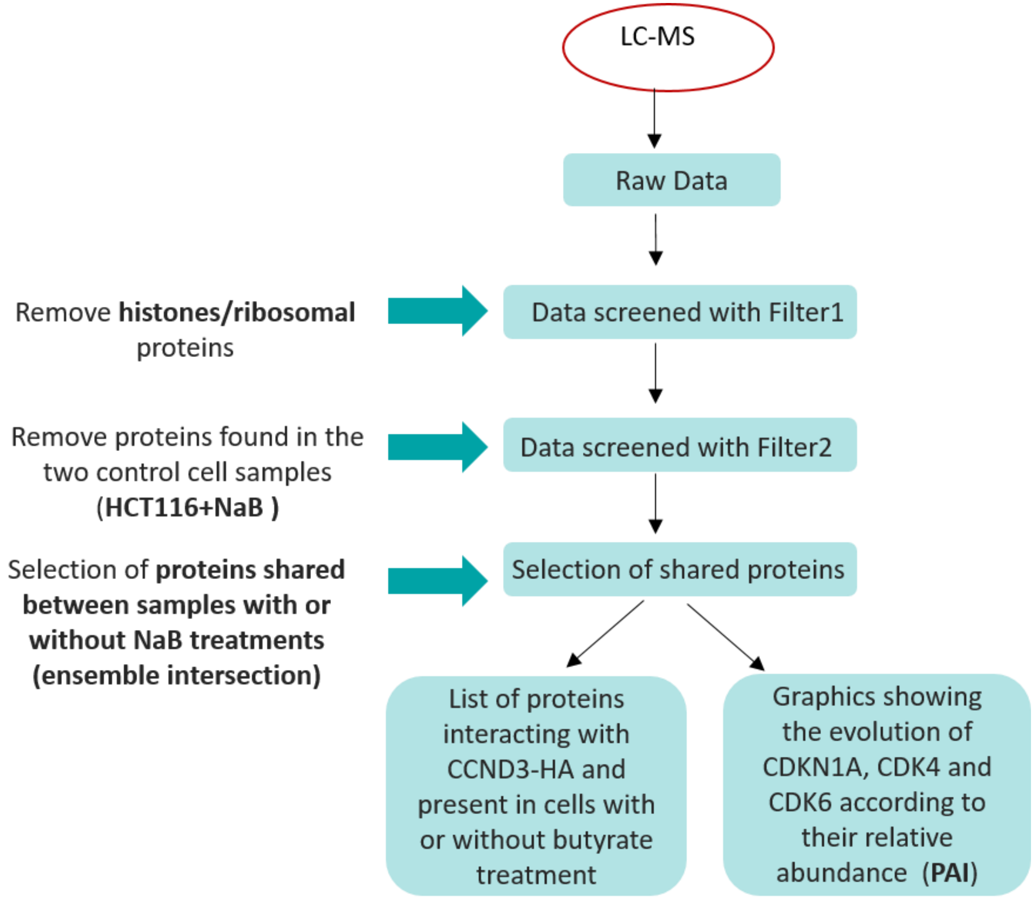

**Fig. S5**: **Venn diagrams showing the selection of the proteins present in samples of P3 and L3 cells cultured in the absence of butyrate**.

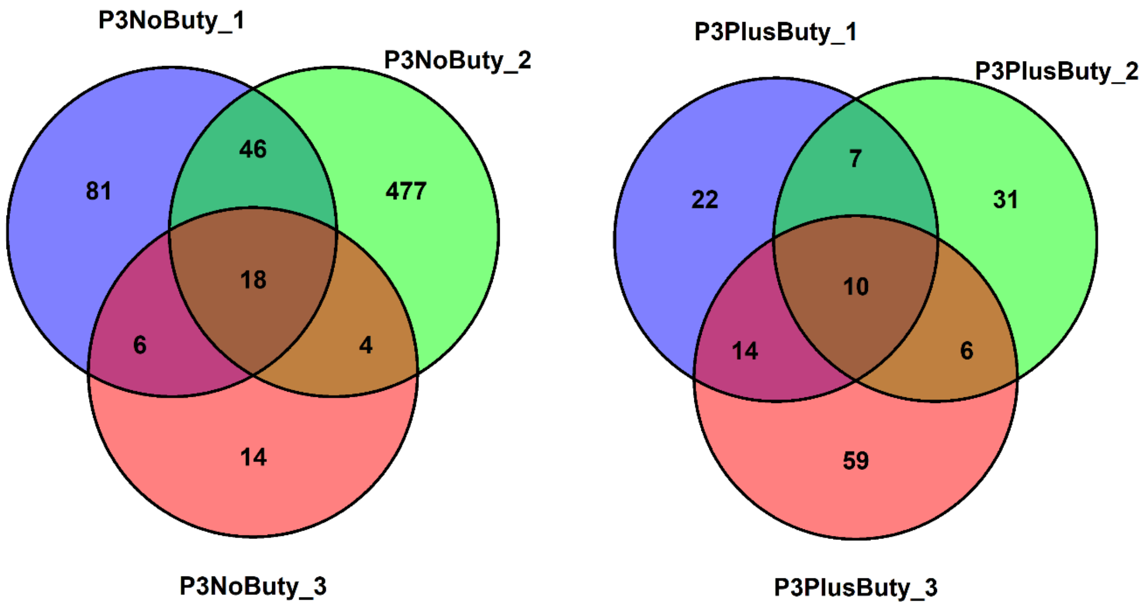

**Fig. S6**: **Venn diagrams showing the selection of the proteins present in samples of P3 and L3 cells cultured in the presence of butyrate**.

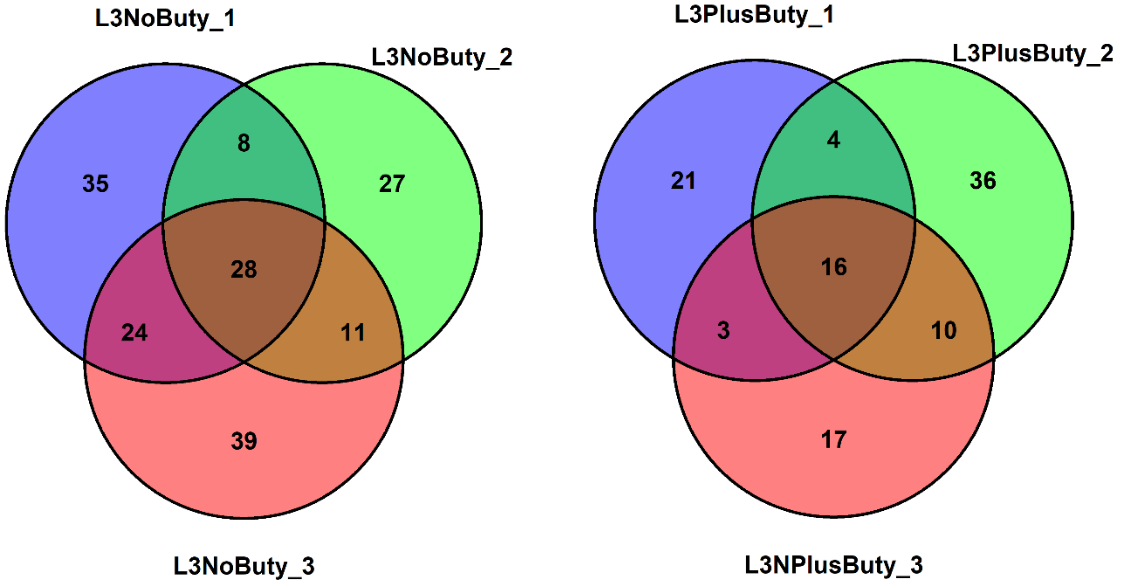

**Fig. S7**: **Venn diagrams showing the number of proteins exclusively present (left) in both samples of L3 and P3 cells** cultured in the absence of butyrate, and (right) in both samples of L3 and P3 cells cultured in the presence of butyrate. In both cases, two proteins were actually CCND3 and CCND3-HA. Therefore, 3 proteins and 4 proteins are interacting with CCND3-HA in cells cultured without or with NaB respectively.

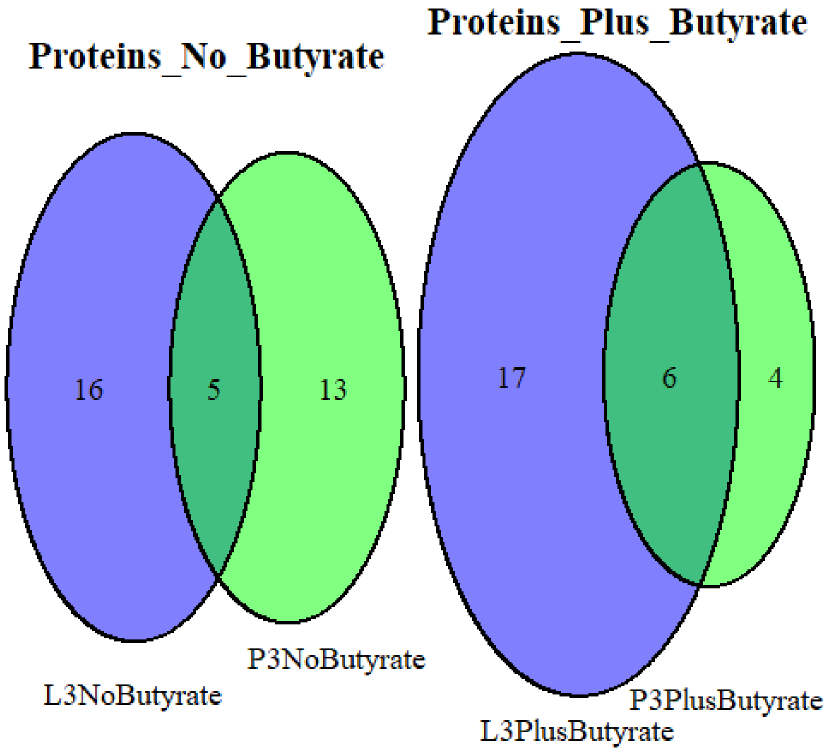


CDKN1A
CDK4
CDK6
CCND3-HA
CCND3

CDKN1A
CDK4
CDK5
CDK6
CCND3-HA
CCND3

### Supplementary data related to the structural analysis and prediction.

**Fig. S8: Multiple sequence alignment of human type D cyclins was performed with CLUSTAL O(1.2.4)** {Sievers, 2011 #16480}. Human CCND3 (P30281) shares 52.25% identity with CCND1 (P24385) and 63.51% identity with CCND2 (P30279). Indicated structures for CCN1 and CCND3 are from PDB entries 2W96 {Day, 2009 #16251} and 3G33 (CCND3-CK4 dimer) respectively Helix Turn Beta

```
sp|P30281|CCND3_HUMAN      ME--LLCCEGTRHAPRAGPDPRLLGDQRVLQSLLRLEERYVPRASYFQCVQREIKPHMRK      58
sp|P24385|CCND1_HUMAN      MEHQLLCCEVE-TIRRAYPDANLLN-DRVLRAMLKAEETCAPSVSYFKCVQKEVLPSMRK      58
sp|P30279|CCND2_HUMAN      ME--LLCHEVD-PVRRAVRDRNLLRDDRVLQNLLTIEERYLPQCSYFKCVQKDIQPYMRR      57
                           **  *** *      **  * .**  :***: :*  **   *  ***:***::: * **:

sp|P30281|CCND3_HUMAN      MLAYWMLEVCEEQRCEEEVFPLAMNYLDRYLSCVPTRKAQLQLLGAVCMLLASKLRETTP      118
sp|P24385|CCND1_HUMAN      IVATWMLEVCEEQKCEEEVFPLAMNYLDRFLSLEPVKKSRLQLLGATCMFVASKMKETIP      118
sp|P30279|CCND2_HUMAN      MVATWMLEVCEEQKCEEEVFPLAMNYLDRFLAGVPTPKSHLQLLGAVCMFLASKLKETSP      117
                           ::* *********:***************:*:  *. *::******.**::***::** *

sp|P30281|CCND3_HUMAN      LTIEKLCIYTDHAVSPRQLRDWEVLVLGKLKWDLAAVIAHDFLAFILHRLSLPRDRQALV      178
sp|P24385|CCND1_HUMAN      LTAEKLCIYTDNSIRPEELLQMELLLVNKLKWNLAAMTPHDFIEHFLSKMPEAEENKQII      178
sp|P30279|CCND2_HUMAN      LTAEKLCIYTDNSIKPQELLEWELVVLGKLKWNLAAVTPHDFIEHILRKLPQQREKLSLI      177
                           ** ********::: *.:* : *::::.****:***:  ***: .:* ::   .:.  ::

sp|P30281|CCND3_HUMAN      KKHAQTFLALCATDYTFAMYPPSMIATGSIGAAVQGLGACSM----SGDELTELLAGITG      234
sp|P24385|CCND1_HUMAN      RKHAQTFVALCATDVKFISNPPSMVAAGSVVAAVQGLNLRSPNNFLSYYRLTRFLSRVIK      238
sp|P30279|CCND2_HUMAN      RKHAQTFIALCATDFKFAMYPPSMIATGSVGAAICGLQQDEEVSSLTCDALTELLAKITN      237
                           :******:****** .*   ****:*:**: **: **        ::   **.:*: :

sp|P30281|CCND3_HUMAN      TEVDCLRACQEQIEAALRESLREASQTSSSPAPKAPRGSSSQGPSQ--TSTPTDVTAIHL      292
sp|P24385|CCND1_HUMAN      CDPDCLRACQEQIEALLESSLRQAQQNMD---PKAAEEEEEEEEEVDLACTPTDVRDVDI      295
sp|P30279|CCND2_HUMAN      TDVDCLKACQEQIEAVLLNSLQQYRQDQRD-GSKSEDELD-------QASTPTDVRDIDL      289
                            : ***:******** * .**::  *       *     ...:  . :.*****   :.:
```

**Fig. S9: In the 6P8H_1 crystallographic structure, the CCND1 sequence is unresolved at its termini**.

```
sp|P24385|CCND1_HUMAN        MEHQLLCCEVETIRRAYPDANLLNDRVLRAMLKAEETCAPSVSYFKCVQKEVLPSMRKIV    60
6P8H_1|Chain                 ------------------DANLLNDRVLRAMLKAEETCAPSVSYFKCVQKEVLPSMRKIV    42
                                               ******************************************

sp|P24385|CCND1_HUMAN        ATWMLEVCEEQKCEEEVFPLAMNYLDRFLSLEPVKKSRLQLLGATCMFVASKMKETIPLT    120
6P8H_1|CCND1(PDB:6P8H)       ATWMLEVCEEQKCEEEVFPLAMNYLDRFLSLEPVKKSRLQLLGATCMFVASKMKETIPLT    102
                             ************************************************************

sp|P24385|CCND1_HUMAN        AEKLCIYTDNSIRPEELLQMELLLVNKLKWNLAAMTPHDFIEHFLSKMPEAEENKQIIRK    180
6P8H_1|CCND1(PDB:6P8H)       AEKLCIYTDNSIRPEELLQMELLLVNKLKWNLAAMTPHDFIEHFLSKMPEAEENKQIIRK    162
                             ************************************************************

sp|P24385|CCND1_HUMAN        HAQTFVALCATDVKFISNPPSMVAAGSVVAAVQGLNLRSPNNFLSYYRLTRFLSRVIKCD    240
6P8H_1|CCND1(PDB:6P8H)       HAQTFVALCATDVKFISNPPSMVAAGSVVAAVQGLNLRSPNNFLSYYRLTRFLSRVIKCD    222
                             ************************************************************

sp|P24385|CCND1_HUMAN        PDCLRACQEQIEALLESSLRQAQQNMDPKAAEEEEEEEEEVDLACTPTDVRDVDI         295
6P8H_1|CCND1(PDB:6P8H)       PDCLRACQEQIEALLESSLRQAQQNMD----------------------------         249
                             ***************************
```

**Fig. S10: Sequence alignment between the whole CCND3 sequence and the partial CCND1 sequence resolved in the in the 6P8H_1 crystallographic structure.** The sequence identity falls to 56.2% % identity outside IDDs (47.9% compared to the whole CCND3 sequence). The difference comes both from the unresolved termini in the crystal structure and the resolved parts, notably the NRLSP sequence that is absent in CCND3.

```
sp|P30281|CCND3_HUMAN      MELLCCEGTRHAPRAGPDPRLLGDQRVLQSLLRLEERYVPRASYFQCVQREIKPHMRKML    60
6P8H_1|Chain               -----------------DA-NLLNDRVLRAMLKAEETCAPSVSYFKCVQKEVLPSMRKIV    42
                                            *   * ::***:::*: **  .* .***:***:*: * ***::

sp|P30281|CCND3_HUMAN      AYWMLEVCEEQRCEEEVFPLAMNYLDRYLSCVPTRKAQLQLLGAVCMLLASKLRETTPLT    120
6P8H_1|Chain               ATWMLEVCEEQKCEEEVFPLAMNYLDRFLSLEPVKKSRLQLLGATCMFVASKMKETIPLT    102
                           * *********:***************:**  *.:*::******.**::***::** ***

sp|P30281|CCND3_HUMAN      IEKLCIYTDHAVSPRQLRDWEVLVLGKLKWDLAAVIAHDFLAFILHRLSLPRDRQALVKK    180
6P8H_1|Chain               AEKLCIYTDNSIRPEELLQMELLLVNKLKWNLAAMTPHDFIEHFLSKMPEAEENKQIIRK    162
                            ********::: *.:* : *:*::.****:***:  ***: .:* ::   .:.: :::*

sp|P30281|CCND3_HUMAN      HAQTFLALCATDYTFAMYPPSMIATGSIGAAVQGLGACSM----SGDELTELLAGITGTE    236
6P8H_1|Chain               HAQTFVALCATDVKFISNPPSMVAAGSVVAAVQGLNLRSPNNFLSYYRLTRFLSRVIKCD    222
                           *****:****** .*   ****:*:**: ******.  *     *  .**.:*: :   :

sp|P30281|CCND3_HUMAN      VDCLRACQEQIEAALRESLREASQTSSSPAPKAPRGSSSQGPSQTSTPTDVTAIHL 292
6P8H_1|Chain               PDCLRACQEQIEALLESSLRQAQQNMD----------------------------- 249
                            ************ *..***:*.*. .
```

**Fig. S11: CDN1A_HUMAN Cyclin-dependent kinase inhibitor** 1. Zn Finger (**small letters in bold**), rrl motif required to bind cyclins (red in bold), region required for binding CDKs (yellow highlight) and IDD (regular small letters) are indicated according to P38936 entry (ProtParam program, Expasy.org)**.**

```
                 *                 *                 *                 *                 *                 *
msepagdvrqnpcgskacrrlfgpvdseqlsrdcdalmagcIQEARERWNFDFVTETPLE
                 *                 *                 *                 *                 *                 *
gdfawervrglglpklylptgprrgrdelgggrrpgtspallqgtaeedhvdlslsctlv
                 *                 *                 *                 *
prsgeqaegspggpgdsqgrkrrqtsmtdfyhskrrlifskrkp
```

**Fig. S12***:* **Per-residue: mean disorder propensities (MDPs), binding-induced folding propensity (ANCHOR2), and liquid-liquid phase separation (LLPS) propensities (FuzDrop)** for (a) CDKN1A, (b) CCND1, (c) CCND3, (d) CDK4 and (e) CDK5, evaluated using Rapid Intrinsic Disorder Analysis Online (RIDAO). Values range from 0 to 1 and a threshold of 0.5 (small-dash line) is used to distinguish ordered residues (MDP < .5) from disordered residues (MDP >=.5) as well as regions that fold upon binding (ANCHOR2 >=.5) from those that do not (ANCHOR2 <.5). Similarly, a threshold of 0.6 (large-dash line) is used to discriminate regions prone to LLPS (FuzDrop >=.6) from those that are not prone to LLPS (FuzDrop < .6). The shaded gray region represents the standard deviation among RIDAO constituent predictors—PONDR VL-XT, PONDR VSL2B, PONDR VL3, PONDR-FIT, IUPred (long), and IUPred (short)—that are used to compute the MDP of each residue.

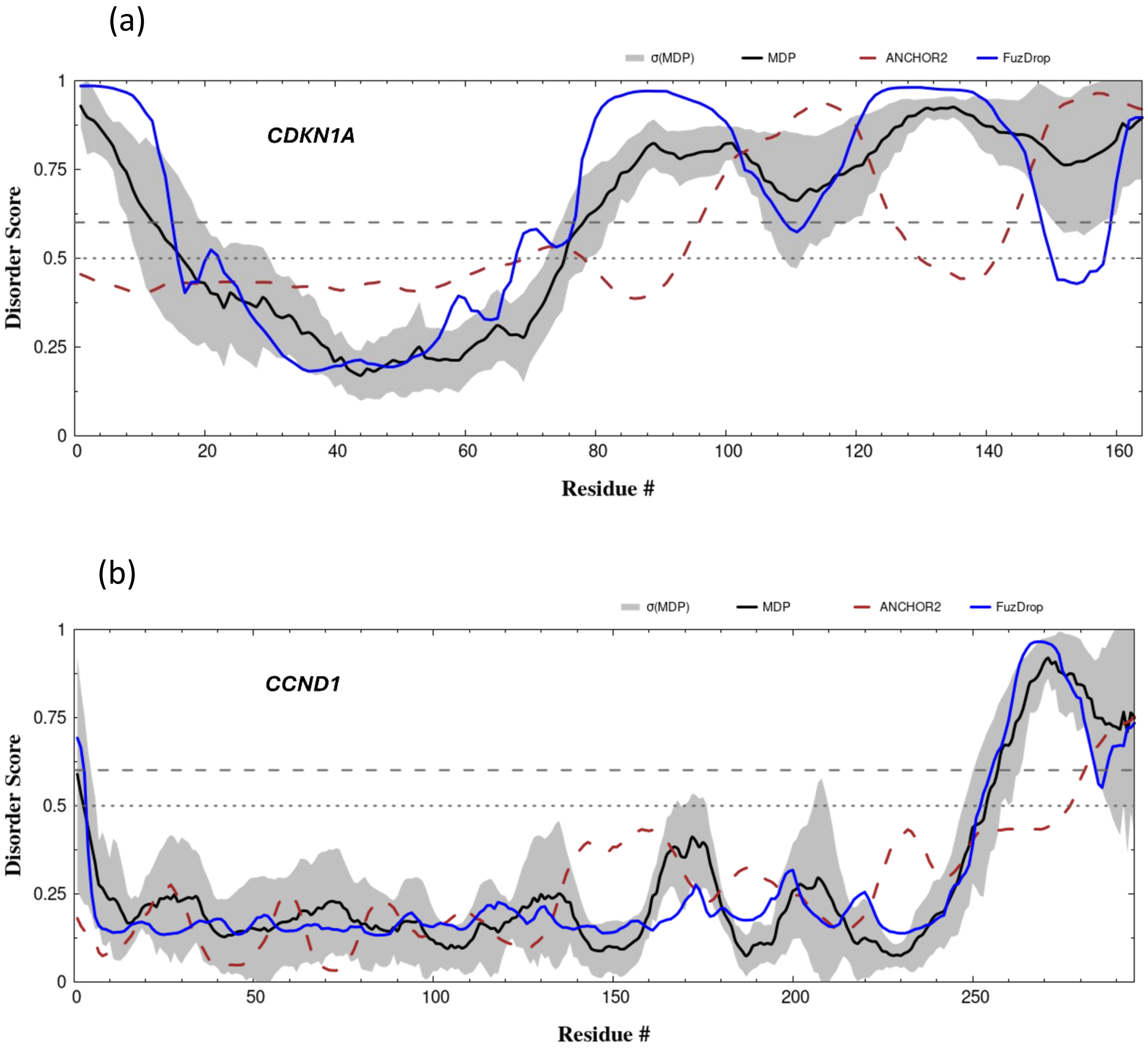
(a)
σ(MDP)
MDP
ANCHOR2
FuzDrop
CDKN1A
Disorder Score
Residue #
(b)
σ(MDP)
MDP
ANCHOR2
FuzDrop
CCND1
Disorder Score
Residue #

(c)

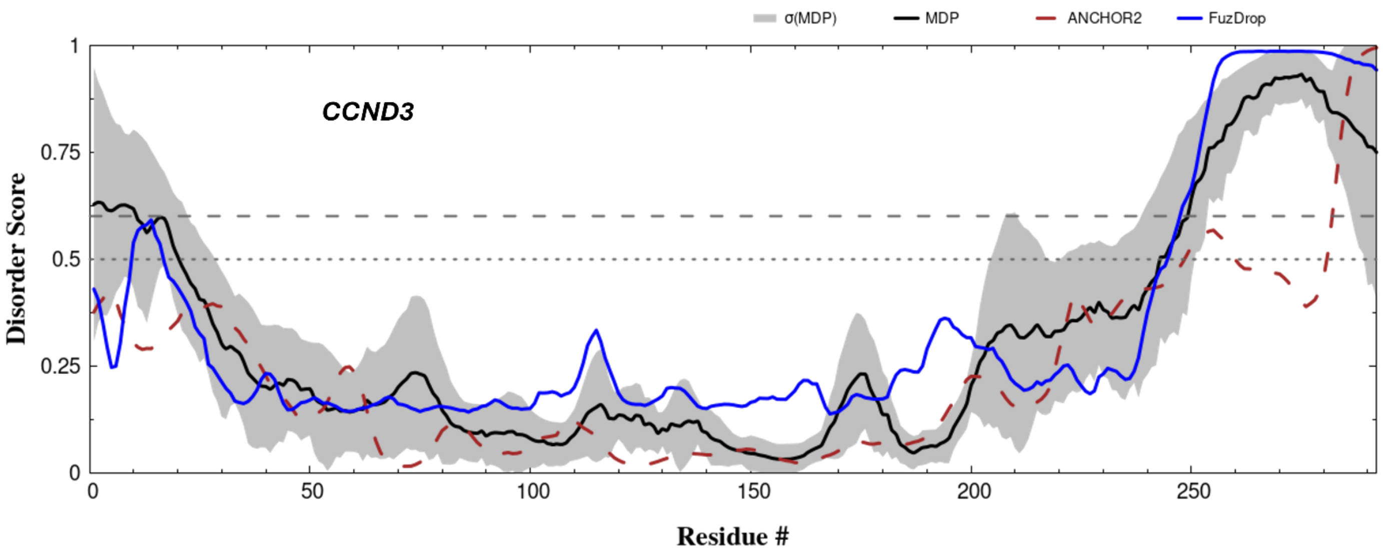
σ(MDP)
MDP
ANCHOR2
FuzDrop
CCND3
Disorder Score
Residue #
1
0.75
0.5
0.25
0
0
50
100
150
200
250

(d)

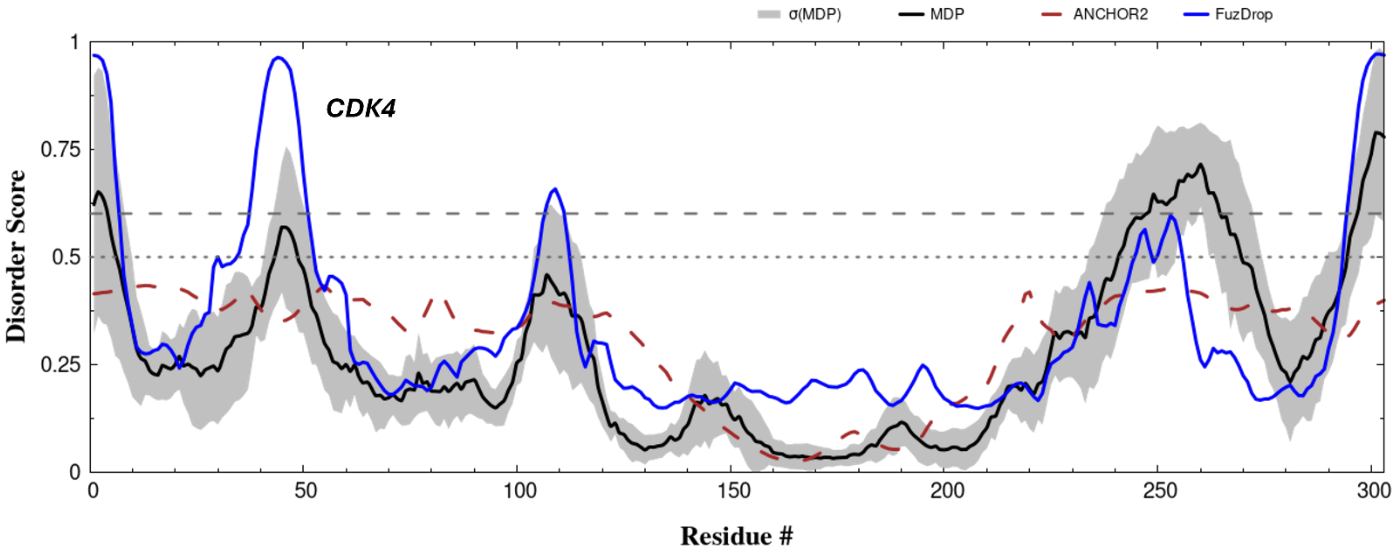
σ(MDP)
MDP
ANCHOR2
FuzDrop
CDK4
Disorder Score
Residue #
0
0.25
0.5
0.75
1
0
50
100
150
200
250
300

(e)

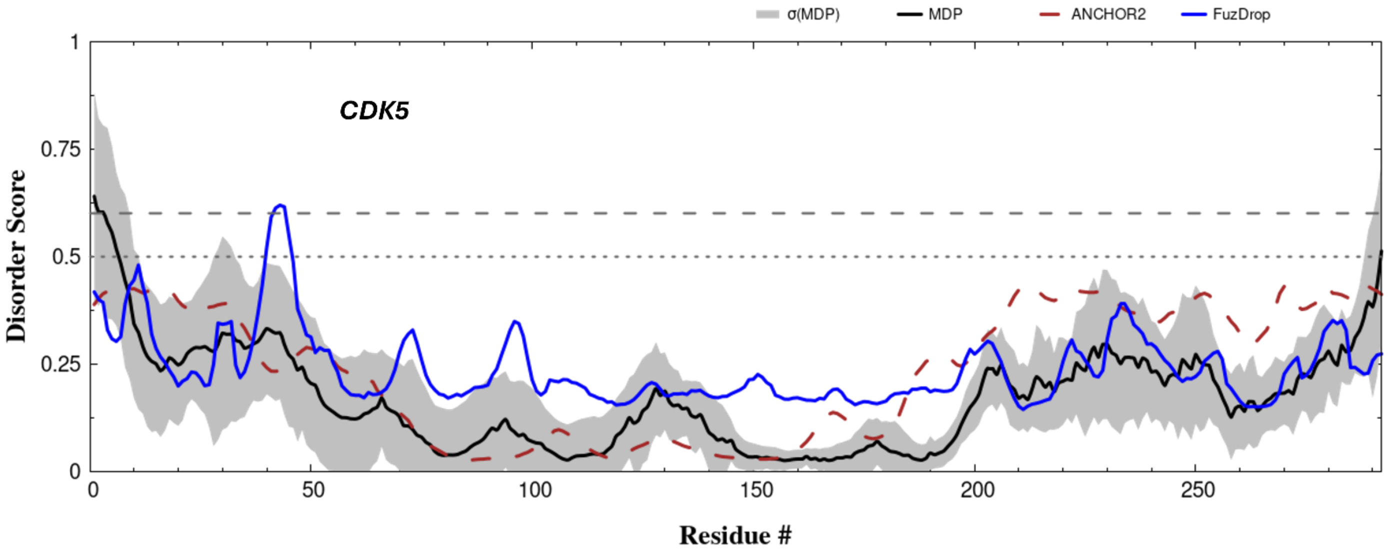
σ(MDP)
MDP
ANCHOR2
FuzDrop
CDK5
Disorder Score
Residue #
0
0.25
0.5
0.75
1
0
50
100
150
200
250
